# CDK19 and CDK8 Mediator kinases drive androgen-independent *in vivo* growth of castration-resistant prostate cancer

**DOI:** 10.1101/2023.08.08.552491

**Authors:** Jing Li, Thomas A. Hilimire, Yueying Liu, Lili Wang, Jiaxin Liang, Balazs Gyorffy, Vitali Sikirzhytski, Hao Ji, Li Zhang, Chen Cheng, Xiaokai Ding, Kendall R. Kerr, Charles E. Dowling, Alexander A. Chumanevich, Gary P. Schools, Chang-uk Lim, Xiaolin Zi, Donald C. Porter, Eugenia V. Broude, Campbell McInnes, George Wilding, Michael B. Lilly, Igor B. Roninson, Mengqian Chen

## Abstract

Castration-resistant prostate cancer (CRPC) remains incurable due to its high plasticity. We found that Mediator kinases CDK8 and CDK19, pleiotropic regulators of transcriptional reprogramming, are differentially affected by androgen, which downregulates CDK8 and upregulates CDK19. Accordingly, expression of CDK8 decreases while CDK19 increases during prostate carcinogenesis, but both CDK19 and CDK8 are upregulated in metastatic CRPC. Genetic inactivation of CDK8 and CDK19 suppresses CRPC tumor growth in castrated male mice and renders CRPC responsive to androgen deprivation. Restoration of active CDK19 or CDK8 kinases reverses this phenotype, indicating that CRPC becomes dependent on Mediator kinase activity for *in vivo* growth under the conditions of androgen deprivation. Selective CDK8/19 inhibitors suppress androgen-independent growth of cell line-based and patient-derived CRPC xenografts, whereas prolonged inhibitor treatment induces tumor regression and even leads to cures. Mediator kinase activity was found to affect tumor and stromal gene expression preferentially in castrated mice, orchestrating castration-induced transcriptional reprogramming. These results warrant the exploration of Mediator kinase inhibitors for CRPC therapy.

## INTRODUCTION

Androgen deprivation therapy (ADT) is the mainstay of treatment of prostate cancer (PCa), the most common cancer and the second leading cause of cancer related mortality in the US men.^1^ However, most patients with aggressive PCa become resistant to ADT and develop castration-resistant prostate cancer (CRPC). Newer CRPC treatments targeting androgen receptor (AR) or androgen production, such as enzalutamide and abiraterone, extend CRPC survival by only 2–8 months, and CRPC remains an incurable disease.^2^ Multiple mechanisms of ADT resistance have been identified, such as increased expression of AR, mutations in its ligand-binding domain, and production of androgen-independent AR splice variants, as well as changes in TP53, RB1 and ETS family.^3^ It has now become apparent that cancer treatment not only selects for genetically altered drug-resistant cells but also induces mass-scale non-genetic adaption, with transcriptional reprogramming leading to drug resistance.^4^ Transcriptional mechanisms of ADT resistance are prominent in PCa, where multiple non-genetic resistance mechanisms can co-exist in the same tumor due to the high heterogeneity of AR expression and other key drivers of PCa growth,^3^ and where the tumor microenvironment acts as another determinant of ADT resistance.^5,6^

CDK8 and CDK19 Mediator kinases are alternative enzymatic components of the kinase module, which binds and regulates the transcriptional Mediator complex. The Mediator kinase module includes, in addition to CDK8 or CDK19, their binding partner Cyclin C (CCNC), as well as MED12/MED12L and MED13/MED13L.^7^ CDK8 and CDK19 paralogs have qualitatively similar effects on protein phosphorylation and transcription but the expression of CDK8 and CDK19 is differentially regulated.^8^ CDK8/19 kinase activities regulate transcription both positively, by potentiating the induction of gene expression by various signals and stressors,^8–11^ and negatively; the latter effect involves post-transcriptional regulation of the Mediator complex proteins by CDK8/19.^8^ Mediator kinases have been identified as broad-spectrum positive regulators of transcriptional reprogramming^8,10,11^ but they also act as negative regulators of chemically induced reprogramming of the cell fate.^12^ Consistently with the effect of Mediator kinases on transcriptional reprogramming, CDK8/19 inhibitors (CDK8/19i) were found to prevent the development of resistance and sometimes overcome the already acquired resistance to different classes of anticancer agents, *in vitro* and *in vivo.*^13–17^

Selective Mediator kinase inhibitors show pronounced single-agent *in vivo* activity in breast cancer^13,15^ and metastatic colon cancer.^18^ The strongest antiproliferative effects of CDK8/19i have been found in acute myeloid leukemia (AML).^19^ In the case of AML, Mediator kinase-regulated genes were associated with super-enhancers (which are enriched in the Mediator complex), and CDK8/19 inhibition hyper-activated such genes. Remarkably, both the inhibition and the upregulation of these genes inhibited AML proliferation, indicating that unbalanced transcription of super-enhancer associated genes was responsible for the anti-leukemic effect.^19^ In addition to their effects on tumor cells, CDK8/19i were also shown to stimulate tumor surveillance by NK cells^20,21^ and effector T-cells.^22^ Although systemic toxicity was reported for two CDK8/19i,^23^ it was subsequently shown to be due to the off-target effects of these compounds,^24^ and several Mediator kinase inhibitors have reached clinical trials in solid tumors and leukemias (clinicaltrials.gov NCT03065010, NCT04021368, NCT05052255, NCT05300438).^25^

Mediator kinases show remarkable clinical correlations in PCa, the only type of cancer marked by high expression of CDK19 in primary tumors.^26^ CDK19 in PCa correlates with Gleason grade, T-stage, Ki67 index, nuclear AR expression and ERG status^27^ and can be used as a marker for the detection of advanced PCa.^28^ Both CDK19 and CDK8 are elevated in metastatic CRPC (mCRPC),^26,27^ and the expression of CDK19^26^ and CDK8^29^ shows negative correlation with disease-free survival. CDK8/19-inhibiting small molecules have been reported to inhibit proliferation of a single PCa cell line,^30^ potentiate the antiproliferative effects of antiandrogens,^31^ and suppress the invasive growth of PCa cells.^26,32^ The function and regulation of CDK8 and CDK19 in PCa are poorly understood, and the potential utility of Mediator kinase inhibitors for PCa treatment remains unclear.

In the present study, we have used genetic modification of CDK19 and CDK8 and selective Mediator kinase inhibitors to investigate the role of Mediator kinase inhibition in the androgen-independent growth of CRPC cell line-based and patient-derived xenograft (PDX) model. We have found that CDK8/19 kinase activity restrains castration-induced transcriptional reprogramming and drives androgen-independent *in vivo* growth of those PCa that have acquired the CRPC phenotype. These results warrant the exploration of selective Mediator kinase inhibitors as a new class of agents for the treatment of the presently incurable mCRPC.

## RESULTS

### Differential effects of androgen signaling on CDK8 and CDK19 expression and upregulation of Mediator kinase module in mCRPC

Analysis of CDK8 and CDK19 gene expression in 4798 normal and 7843 tumor clinical tissue samples from 16 different organs (Fig. 1A) was conducted using the RNA-Seq data in the TNMplot database.^33^ This analysis revealed that CDK19 is expressed in androgen-dependent organs, prostate and testes, higher than in any other normal tissues and that CDK19 expression further increases in PCa, reaching higher levels than in any other cancers; in contrast, CDK19 greatly decreases during testicular carcinogenesis. On the other hand, CDK8 is expressed at an intermediate level in normal prostate and at a high level in testes but it is downregulated in both PCa and testicular cancers relative to their normal tissue counterparts (Fig. 1A). Preferential elevation of CDK19 RNA expression in PCa is also seen among the cell lines in the DepMap.org database. Among the top 1% of cell lines with the highest CDK19 expression (14 out of 1450), 4 (29%) belong to the prostate lineage, which constitutes < 1% of all cell lines (12 out of 1450). Among these, 7 cell lines representing prostate adenocarcinoma show a strong correlation between CDK19 and AR expression (Fig. 1B). The association between CDK19 and AR expression was confirmed at the protein level by Western blot analysis in Fig. 1C, which compares CDK8, CDK19 and AR (including full-length AR (AR-FL) and its splice variants (AR-Vs)) in different prostate cancer cell lines (293 cells and 293 with CRISPR-mediated knockout of both CDK8 and CDK19 (293-dKO)^34^ were used as reference controls). All the PCa cell lines express less CDK8 than 293, whereas CDK19 expression is strongly increased in AR-positive but not in AR-negative PCa (Fig. 1C).

**Figure 1.**
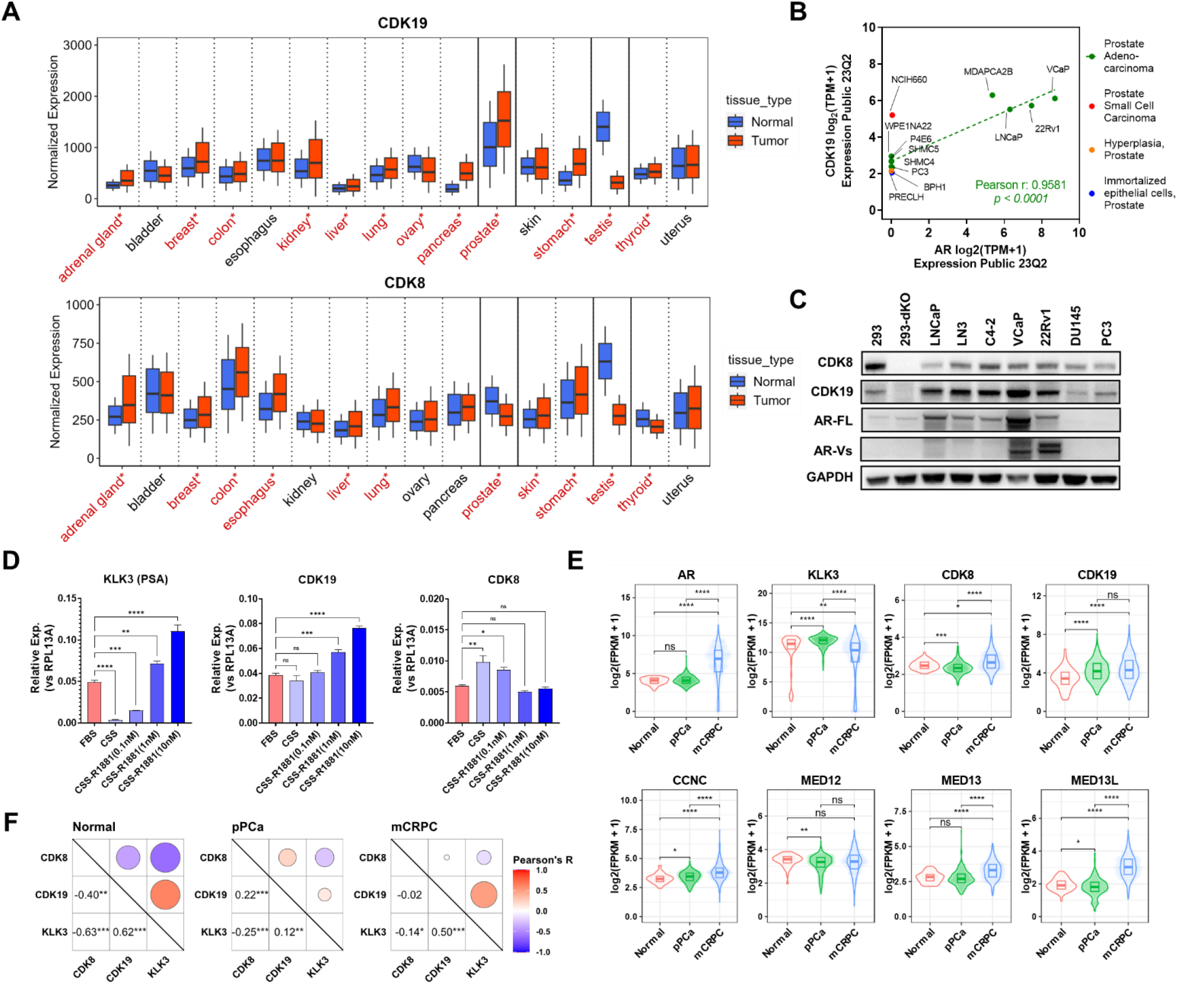
Expression of CDK8 and CDK19 in PCa. **(A)** CDK19 and CDK8 RNA expression in normal and tumor tissues. Significant differences between normal and tumor tissues (Mann-Whitney U test) are marked in red and with asterisks. **(B)** RNA expression of CDK19 and AR in 12 prostate-lineage cell lines from the DepMap database. Pearson correlation analysis was performed for prostate adenocarcinomas (green dots). **(C)** Western blot analysis of CDK8, CDK19, AR and GAPDH (control) in different prostate cancer cell lines and in 293 cells and their CDK8/19 double knockout (dKO) derivative. **(D)** qPCR analysis of KLK3 (PSA), CDK19, and CDK8 RNA in LNCaP cells growing in androgen-containing (FBS) or androgen-deprived (CSS) media for 3 days and treated with or without R1881 androgen at indicated concentrations for 24 hours. **(E)** RNA expression of AR, KLK3, and Mediator-associated CDK module subunits in normal prostate (n = 52), primary PCa (n = 502) and metastatic CRPC (n = 266) tissues. **(F)** Correlation analysis among CDK8, CDK19 and KLK3 expression in normal prostate, primary PCa and mCRPC clinical samples. Asterisks in (D-F) mark p-values: 0 < **** < 0.0001 < *** < 0.001 < ** < 0.01 < * < 0.05. ns, not significant.

We have analyzed the effects of androgen signaling on CDK8 and CDK19 RNA expression in androgen-responsive LNCaP cells. Cells were transferred from medium with androgen-containing FBS to medium with androgen-depleted CSS (charcoal-stripped serum) for 48 hours, followed by the addition of R1881 androgen at 0.1, 1 and 10 nM for 24 hours. qPCR analysis (Fig. 1D) showed that the expression of KLK3 (PSA) gene driven by canonical AR signaling was abrogated by androgen depletion but induced by androgen addition, in a concentration-dependent manner. CDK19 expression was unaffected by androgen depletion but upregulated by androgen addition, whereas CDK8 expression was increased by androgen depletion but decreased to the basal level by androgen addition (Fig. 1D). Hence, androgen signaling positively regulates CDK19 and negatively regulates CDK8 expression.

We have compared androgen signaling and RNA expression of the Mediator kinase module subunits using RNA-Seq data from the TCGA and cBioPortal databases, for normal prostate tissues, primary^35^ and metastatic PCa (the latter samples come from patients that have failed ADT and therefore can be classified as mCRPC^36^). The expression of AR and KLK3 was elevated in primary tumors relative to normal prostate tissues, reflecting carcinogenesis-associated increase in canonical AR signaling. On the other hand, mCRPC showed a strong increase in AR but a decrease in KLK3 relative to primary tumors, indicating a change from canonical AR-driven transcriptional signaling (Fig. 1E). CDK8 was downregulated in primary tumors, consistently with its negative regulation by androgen in LNCaP cells, but CDK8 was strongly increased in mCRPC. In contrast, CDK19, which is positively regulated by androgen signaling, was strongly increased in primary PCa and further increased in mCRPC (although the increase of CDK19 from primary PCa to mCRPC was not statistically significant here, this was previously documented in other studies^28,37^). Expression of all the other Mediator kinase module components: CCNC, MED12, MED13, MED13L (MED12L isoform was expressed at a very low level in PCa) increased from normal to primary to metastatic PCa (although the increases from normal to primary tumors for MED13 and from primary to metastatic PCa for MED12 were not statistically significant) (Fig. 1E). We have determined pairwise Pearson correlation coefficients between the expression of CDK8, CDK19 and KLK3 (used as a marker of canonical androgen signaling) among normal, primary and metastatic prostate samples (Fig. 1F). In agreement with the effects of androgen in LNCaP (Fig. 1D), KLK3 expression was positively correlated with CDK19 and negatively correlated with CDK8 in all three sets of tissues (Fig. 1F).

These results suggest that downregulation of CDK8 and upregulation of CDK19 in primary PCa reflect transcriptional effects of elevated androgen signaling, and that altered AR-mediated transcription in mCRPC abrogates the negative regulation of CDK8 expression.

### Effects of CDK8/19 inhibition on androgen-responsive gene expression and cell growth of PCa cells with different degrees of androgen independence

Since Mediator kinases modulate signal-induced transcription, we analyzed the effects of CDK8/19 inhibition on androgen-regulated transcriptional signaling in PCa cells. We have carried out RNA-Seq analysis of androgen-depleted LNCaP cells that were untreated or treated with two chemically unrelated selective CDK8/19 kinase inhibitors (CDK8/19i) Senexin B^13^ at 2 μM or SNX631^15^ at 0.5 μM, with or without R1881 androgen stimulation for 24 or 72 hrs. Differentially Expressed Genes (DEGs) were selected using False Discovery Rate (FDR) < 0.01 and Fold Change (FC) >1.5 as cutoff criteria. DEGs affected by CDK8/19i in androgen-depleted or androgen-stimulated cells are listed in Table S1. Volcano plots in Fig. S1A and Venn diagrams in Fig. S1B show that Mediator kinase inhibitors affected 2-3% of all the genes with expression levels greater than 1 count per million (cpm), while 16-17% of the expressed genes (cpm >1) were impacted by the addition of R1881. Remarkably, the fraction of androgen-affected genes among CDK8/19i-affected DEGs increased to 45-70%, with a greater increase under androgen-deprived conditions.

Fig. 2A shows that Senexin B and SNX631 both potentiated and counteracted the effects of androgen on gene expression. In particular, the induction of three of the most strongly androgen-inducible genes (KLK2, KLK3 (PSA) and CHRNA2) was decreased by the Mediator kinase inhibitors, whereas the expression of UGT2B family genes was inhibited by androgen addition and further inhibited by CDK8/19i (Fig. S1C). Fig. S1D shows the effects of androgen and CDK8/19i on the expression of AR and MYC, key regulators of PCa growth. MYC expression was not significantly affected by androgen, but it was upregulated by both CDK8/19i, in FBS and in CSS, whereas AR expression was mildly downregulated by both androgen and CDK8/19i.

**Figure 2.**
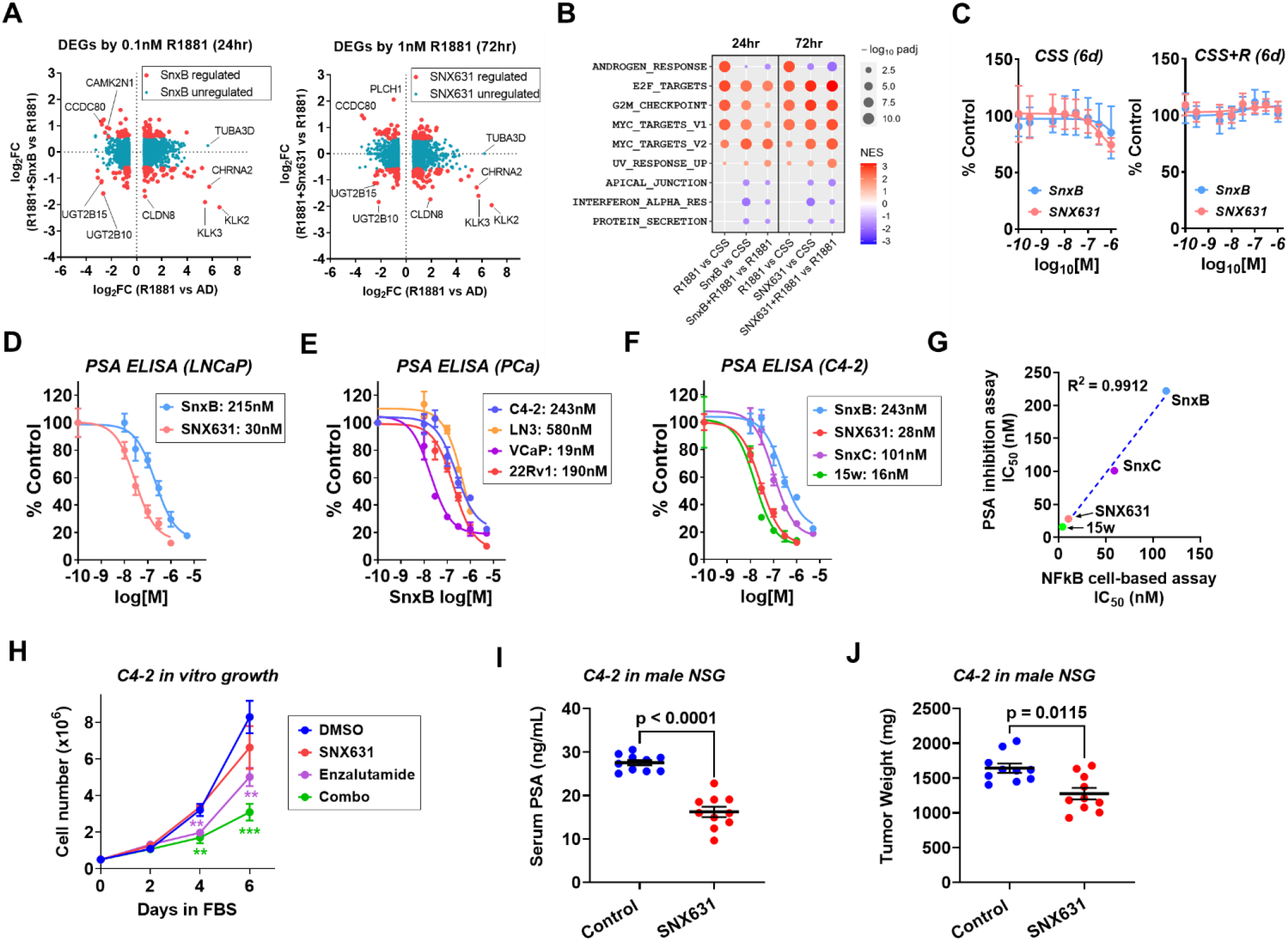
Effects of Mediator kinase inhibition in androgen-responsive PCa cells. **(A)** Effects of treatment with 2 µM Senexin B (SnxB) or 500 nM SNX631 on the androgen-regulated genes in LNCaP cells. **(B)** Hallmark pathways (GSEA) affected by Mediator kinase inhibition in LNCaP cells (CDK8/19i and R1881 concentrations shown in A). **(C)** Effects of Senexin B and SNX631 on the growth of LNCaP cells in CSS media with or without 100 pM R1881, measured by the Sulforhodamine B (SRB) assay. **(D)** Effects of Senexin B and SNX631 treatment (4 days) on PSA protein expression (ELISA) in conditioned media of LNCaP cells in FBS media. **(E)** Effects of Senexin B treatment on PSA expression in conditioned media of different AR-positive PCa cell lines. **(F)** Effects of different CDK8/19i on PSA expression in conditioned media of C4-2 cells. **(G)** Correlation of IC50 values of different CDK8/19i based on PSA ELISA in C4-2 cells and NFκB reporter assay in 293 cells. **(H)** Effects of SNX631 (500 nM) and enzalutamide (5 µM), individually or in combination, on the growth of C4-2 cells in FBS media. (**, p<0.01; ***, p<0.001). **(I,J)** Serum PSA **(I)** and final tumor weights **(J)** of C4-2 xenografts grown in intact male NSG mice, treated with SNX631 (25 mg/kg, b.i.d.) or vehicle for 14 days.

Gene Set Enrichment Analysis (GSEA)^38^ of 50 hallmark pathways identified nine pathways that were significantly affected by Mediator kinase inhibition in LNCaP cells, with or without androgen stimulation. Among these, several pathways associated with cell proliferation (MYC and E2F targets; G2/M) and UV response were upregulated both by androgen and by CDK8/19i (Fig. 2B), suggesting cooperative effects; only the androgen response pathway showed the opposite effects of androgen and CDK8/19i (Fig. S1E).

To determine whether the transcriptomic effects of CDK8/19i are associated with an effect on cell proliferation in the absence or in the presence of androgen, we evaluated the effects of a 6-day treatment with different concentrations of Senexin B and SNX631 on LNCaP cell growth in CSS, with or without the addition of R1881. Despite the upregulation of MYC and proliferation-associated pathways, CDK8/19i mildly inhibited cell growth in androgen-deprived (CSS) media, but it had no significant effect on the cell number when androgen was added (Fig. 2C). Hence, transcriptomic effects of Mediator kinase activity in androgen responsive PCa cells do not translate into major effects on *in vitro* cell proliferation.

KLK3 (PSA) was one of the genes most strongly induced by androgen and downregulated by CDK8/19i in RNA-Seq analysis of LNCaP cells (Fig. 2A). This result was validated at the protein level by measuring secreted PSA in the conditioned media from LNCaP cells treated with different concentrations of Senexin B or SNX631 in androgen containing FBS media (Fig. 2D). Senexin B also suppressed PSA levels in the supernatant of four AR-positive PCa cell lines with constitutively activated androgen signaling, including C4-2 and LN3 derivatives of LNCaP, as well as VCaP, and 22Rv1 (Fig. 2E), indicating the generality of this effect of CDK8/19 on canonical AR signaling. Fig. 2F compares the effects of 4 different CDK8/19i, including Senexin B, SNX631, Senexin C,^25^ and 15w,^39^ on PSA expression in C4-2 cells. The IC_50_ values for PSA inhibition were perfectly correlated (R^2^ = 0.99) with the IC_50_ values for all three CDK8/19i in a standardized cell-based assay ^25,34^ (Fig. 2G), confirming that the effect was mediated by CDK8/19.

We have investigated the effect of SNX631, alone and in combination with AR antagonist enzalutamide, on *in vitro* growth of LNCaP-derived C4-2 cells, which exhibit partial androgen independence but remain androgen-responsive.^40,41^ The effect of the CDK8/19i alone on C4-2 cell proliferation was insignificant, whereas enzalutamide alone had moderate effect (Fig. 2H). The addition of the CDK8/19i, however, augmented the effect of enzalutamide (Fig. 2H), indicating that Mediator kinase activity preferentially affects C4-2 cell growth when androgen signaling is suppressed. We further examined the *in vivo* effects of SNX631 treatment on tumor growth and serum PSA production by C4-2 xenografts in intact male NCG mice. After 11 days of treatment, the CDK8/19i strongly decreased serum PSA levels (Fig. 2I) and moderately but significantly inhibited tumor growth based on final tumor weights (Fig. 2J). C4-2 cells, however, showed a poor tumor take rate in castrated mice in our hands, and therefore *in vivo* effects of Mediator kinase inhibition in this model could not be evaluated under the conditions of androgen deprivation.

### Effects of CDK8 and CDK19 inhibition on in vitro gene expression and proliferation of 22Rv1 CRPC cells

To study the role of Mediator kinases in CRPC, we have used 22Rv1 cells,^42^ a widely used CRPC model that expresses both a mutated version of FL-AR (AR*^ex^*^3d^*^up^*) and its androgen-independent splice variant AR-V7^43,44^ (Fig. 1C). 22Rv1 cells express 4.5 times more CDK19 than CDK8 at the protein level.^8^ The role of CDK8/19 in 22Rv1 was analyzed using both the CDK8/19i SNX631 and genetic modifications of CDK19 and CDK8. As shown in Fig. 3A, 22Rv1 cells with a double knockout of CDK8 and CDK19 (Rv1-dKO) were previously created via CRISPR/Cas9,^8^ and we have now generated Rv1-dKO derivatives that express wild-type CDK19 (dKO-19), its kinase-inactive D173A mutant^45^ (dKO-19M) or insert-free lentiviral vector (dKO-V), as well as dKO-V derivatives expressing wild-type CDK8 (dKO-8) or its kinase-inactive D173A mutant (dKO-8M).^46^ We have also transduced parental 22Rv1 (Rv1-WT) with a lentivirus expressing firefly luciferase, yielding the derivative Rv1-Luc; Rv1-WT and Rv1-Luc were compared in various assays to account for the inherent phenotypic variability of 22Rv1 cells. Fig. 3B depicts western blot analysis of CDK8 and CDK19 expression in different derivatives and the effects of 6-hr treatment with CDK8/19i SNX631 on S727 phosphorylation of STAT1, a well-known substrate of Mediator kinases.^47^ SNX631 decreased STAT1 S727 phosphorylation in cells expressing WT CDK8 and/or CDK19 but not in dKO-Vec cells or in cells expressing the mutant Mediator kinases, confirming that the mutations have abrogated the kinase activity (Fig. 3B).

**Figure 3.**
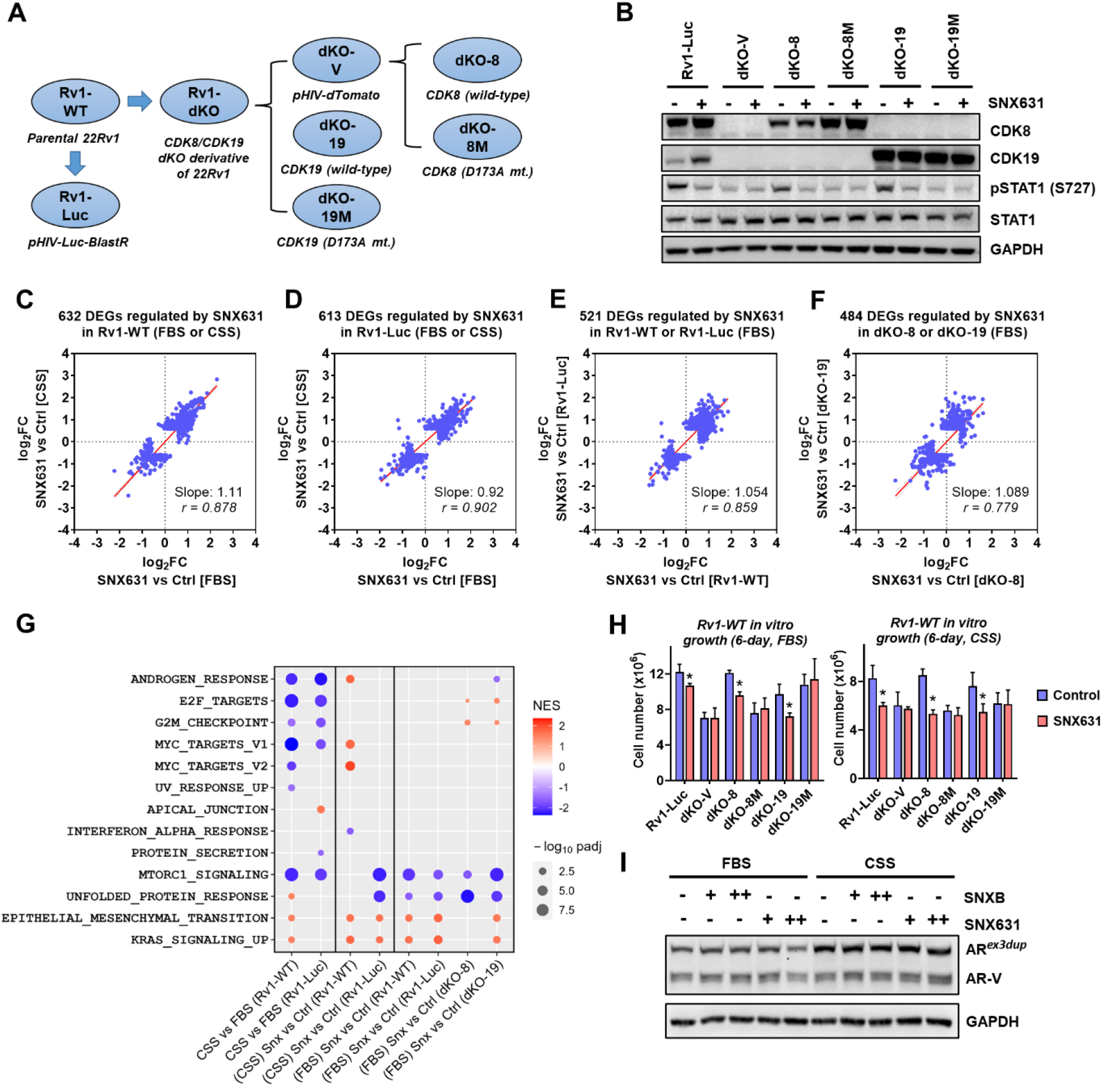
Effects of Mediator kinase inhibition in 22Rv1 derivatives *in vitro*. **(A)** Scheme of generating CDK8/19 double-knockout (dKO) and reconstitution derivatives in 22Rv1 cells. **(B)** Western blot analysis of 22Rv1 derivatives, untreated or treated with 500 nM SNX631 for 6 hours, for the indicated proteins. **(C, D)** Comparison of the effects of Mediator kinase inhibition on SNX631-affected DEGs between androgen-containing (FBS) and androgen-deprived (CSS) conditions in Rv1-WT **(C)** and RV1-Luc **(D)** cells. **(E)** Comparison of the effects of Mediator kinase inhibition on SNX631-affected DEGs between Rv1-WT and Rv1-Luc cells (in FBS media). **(F)** Comparison of the effects of Mediator kinase inhibition on SNX631-affected DEGs between dKO-8 and dKO-19 cells (in FBS media). **(G)** Effects of Mediator kinase inhibition and androgen depletion on the affected hallmark pathways (GSEA) in 22Rv1 derivatives. **(H)** Effects of SNX631 on cell growth of 22Rv1 derivatives in FBS or CSS media. **(I)** Western blot analysis of the effects of CDK8/19i treatment (1 μM SnxB or 500 nM SNX631, 24 hours) on AR in 22Rv1 cells in FBS or CSS media.

We have used RNA-Seq to evaluate the effects of CDK8/19i SNX631 on gene expression in Rv1-WT and Rv1-Luc in androgen-containing (FBS) and androgen-depleted (CSS) media, and in dKO-19, dKO-19M, dKO-8, and dKO-8M derivatives (in FBS). DEGs affected by SNX631 in any derivatives or culture conditions (FC >1.5, FDR < 0.01 cutoffs) are listed in Table S2. Volcano plots in Fig. S2A,B show that growth in androgen-depleted media has affected an order of magnitude more DEGs in Rv1-WT than in RV1-Luc cells, indicating that Rv1-Luc were less androgen-responsive, reflecting the inherent heterogeneity of androgen responsiveness in PCa cell lines.^3,48^ On the other hand, the effects of SNX631 on gene expression in FBS and CSS were similar, both in 22Rv1-WT and in 22Rv1-Luc (Fig. 3C-E). Cell growth in androgen-deprived media affected similar DEG numbers with and without SNX631 treatment (Fig. S2A,B). SNX631 affected very few genes in Mediator kinase-mutated dKO-8M and dKO-19M derivatives, indicating the target selectivity of this CDK8/19i (Fig. S2C,D). SNX631 had similar effects on gene expression in dKO-8 and dKO-19 (Fig. 3F), indicating that transcriptomic effects of CDK8 and CDK19 in 22Rv1 cells are qualitatively similar.

GSEA of the hallmark pathways revealed that androgen deprivation (CSS media) downregulated the androgen response pathway as well as cell proliferation-related pathways (MYC, E2F, and G2M) in Rv1-WT and Rv1-Luc cells, as it did in LNCaP. However, in contrast to LNCaP (Fig. 2B), Mediator kinase inhibition did not upregulate the proliferation-related pathways in most 22Rv1 models (Fig. 3G). Treatment with SNX631 led to a significant downregulation of the unfolded protein response (UPR) and mTORC1 pathways (Fig. 3G). Fig. S2E shows the heatmap of 33 genes that were regulated by CDK8/19 across all the 22Rv1 derivatives and only 4 of these (BASP1, SEMA3C, ARHGAP20, CHAC1) were also regulated by SNX631 in LNCaP cells (Fig. S2F). Hence, CDK8/19 has different transcriptomic effects on PCa cell lines.

The effects of CDK8/19 inhibition on *in vitro* growth of 22Rv1 derivatives were tested in androgen-containing (FBS) and in androgen-deprived (CSS) media via a 6-day assay. SNX631 treatment produced a moderate but significant inhibition of cell proliferation in both FBS and CSS in 22Rv1 derivatives expressing WT CDK8 and/or CDK19 (Rv1-Luc, dKO-8, dKO-19) but not in Mediator kinase-inactive dKO-V, dKO-8M or dKO-19M cells (Fig. 3H). This growth inhibition was not associated with the effects on MYC, which was variably affected by CDK8/19 inhibition in different derivatives (Fig. S2G). AR was unaffected by CDK8/19 inhibition at the RNA (Fig. S2G) or protein levels (Fig. 3I). Hence, CDK8 and CDK19 positively regulate 22Rv1 cell growth *in vitro*, irrespective of androgen supplementation and without altering AR expression.

### Selective effects of CDK8/19 on in vivo growth of 22Rv1 xenografts under the conditions of androgen deprivation

We have investigated the effects of CDK8 and CDK19 expression and kinase activity on *in vivo* growth of 22Rv1 xenografts, with or without androgen deprivation, by monitoring tumor growth of 22Rv1 derivatives in intact or castrated male NSG mice (Fig. 4A). Rv1-WT and Rv1-Luc xenografts grew in both intact and castrated mice; tumor growth in castrated mice was slower for Rv1-WT but not for Rv1-Luc, in agreement with the lesser effect of androgen depletion on gene expression in Rv1-Luc (Fig. S2A, B). CDK8/19 knockout derivatives Rv1-dKO and dKO-V grew in intact mice, but the growth of these tumors in castrated mice was strongly suppressed (Fig. 4A). Re-expression of CDK8 or CDK19 in dKO-8 and dKO-19 derivatives reproduced the phenotype of the Rv1-WT parent, as they grew in both intact and, at a slower rate, in castrated mice. However, dKO-19M and dKO-8M derivatives expressing inactive Mediator kinase mutants formed measurable tumors but were unable to grow in castrated mice (Fig. 4A). These results indicate that the kinase activity of CDK19 or CDK8 is selectively required for *in vivo* growth of 22Rv1 xenografts under the conditions of androgen deprivation.

**Figure 4.**
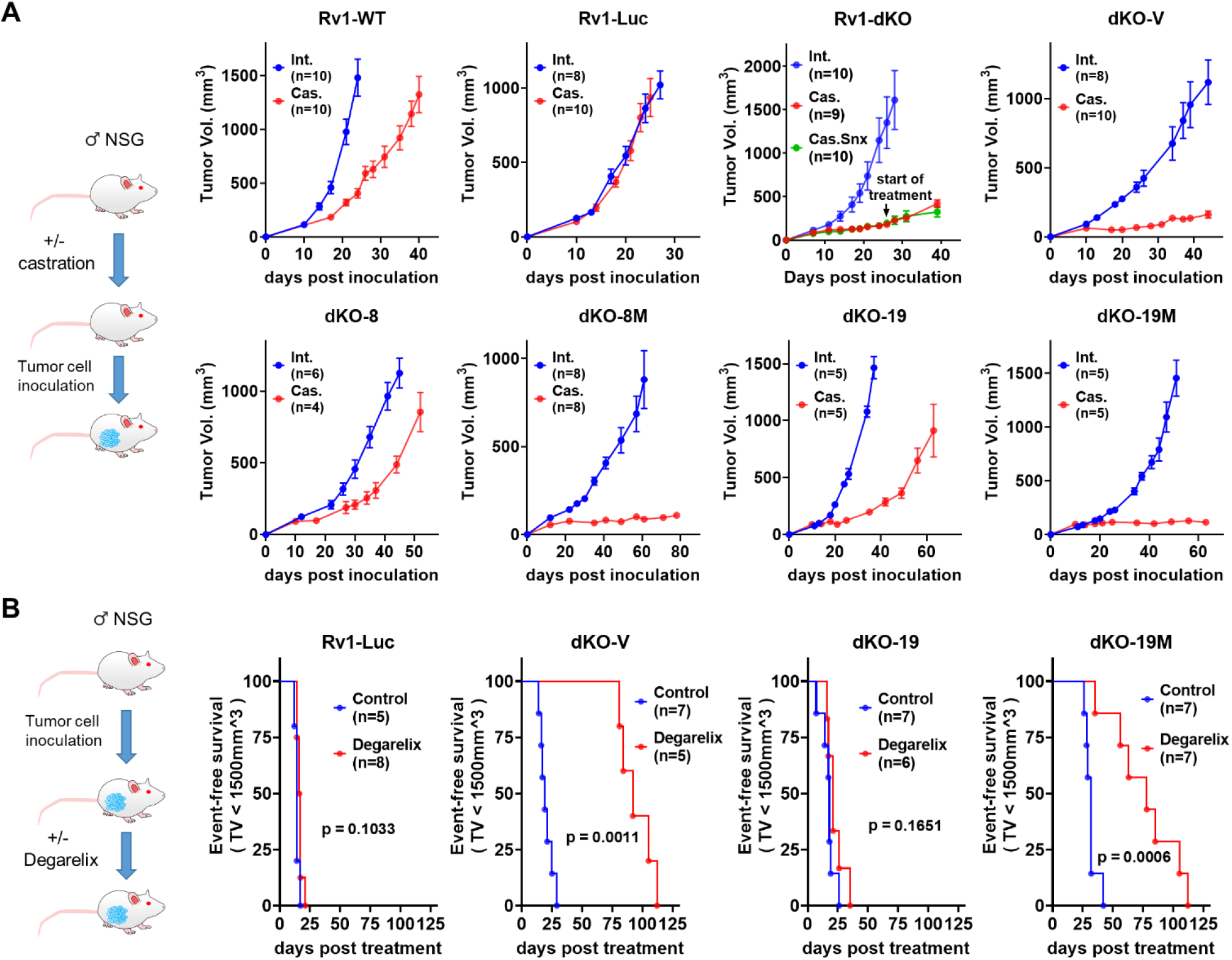
Effects of Mediator kinase mutagenesis on 22Rv1 *in vivo* growth. **(A)** Scheme of the study and xenograft growth curves of the indicated 22Rv1 derivatives in intact and castrated NSG mice. 2×10^6^ tumor cells in 50% Matrigel were subcutaneously (s.c.) implanted into the right flank. Castration was carried out surgically for RV1-WT and Rv1-dKO and chemically (10 mg/kg degarelix, s.c., every 30 days) for Rv1-Luc and dKO derivative studies. Rv1-dKO study also includes an arm where mice were treated with SNX631 (via medicated food, at 500 ppm) starting on day 26 after implantation. **(B)** Scheme of the study and KM plots of the effects of degarelix on event-free survival of NSG mice bearing the indicated 22Rv1 derivatives (event defined as tumor volume exceeding 1.5 cm^3^). Treatment was started when tumor size reached 100-200 mm^3^.

To assess how Mediator kinase activity affects the response of established 22Rv1 tumors to ADT, we have inoculated Rv1-Luc, dKO-V, dKO-19 and dKO-19M derivatives into intact male mice. When the tumors reached 150-200 mm^3^ size, mice were either untreated or treated with degarelix, a gonadotrophin-releasing hormone antagonist that suppresses testosterone production in the body. The effect of degarelix on tumor growth was measured by event-free survival analysis, with the event defined as tumors reaching 1.5 cm^3^ volume. Degarelix treatment did not slow down the growth of Rv1-Luc or dKO-19 tumors that express functional Mediator kinases but drastically inhibited the growth of dKO-V and dKO-19M derivatives that are deficient in Mediator kinase activity (Fig. 4B). Hence, Mediator kinase inhibition restores the response to androgen deprivation in 22Rv1 CRPC tumors.

### Systemic CDK8/19 inhibitor treatment suppresses androgen-independent in vivo growth and produces cures in 22Rv1 xenografts

The effects of systemic *in vivo* treatment with CDK8/19i SNX631 on 22Rv1 tumors growing in intact or castrated male NSG mice were analyzed as diagrammed in Fig. 5A. Similarly to Mediator kinase mutagenesis, SNX631 had little effect on Rv1-Luc (Fig. 5B) or Rv1-WT (Fig. 5C) xenograft growth in intact mice but strongly inhibited tumor growth of both cell lines in castrated mice, as indicated by the effects on tumor volumes and final tumor weights (Fig. 5B,C). SNX631 treatment had no detrimental effect on the body weights in intact or castrated mice compared to vehicle groups (Fig. 5D). Machine learning based histological analysis showed that tumor suppression by SNX631 in castrated mice was associated with decreased cell proliferation and increased necrosis (Fig. 5E,F). SNX631 treatment had no effect on CDK8/19-deficient Rv1-dKO tumors (Fig. 4A), confirming that the inhibitor’s effect was mediated by CDK8/19.

**Figure 5.**
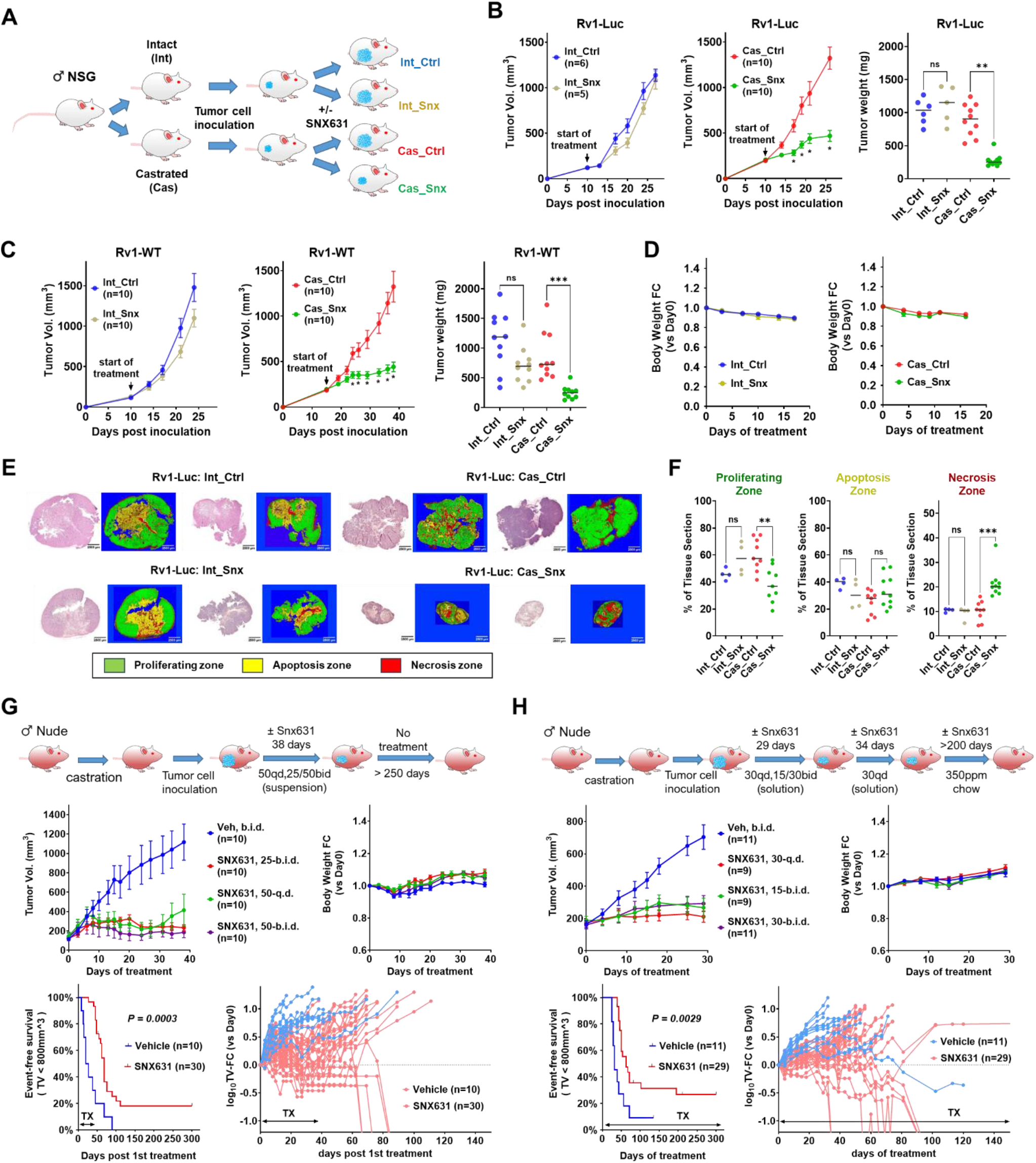
Effects of CDK8/19 inhibitor on 22Rv1 *in vivo* growth. **(A)** Scheme of the study of the effects of SNX631 on 22Rv1 growth in intact and castrated NSG mice. **(B)** Tumor growth curves and final tumor weights of Rv1-Luc xenografts growing in intact or castrated mice, receiving the control or SNX631-medicated diet (500 ppm). **(C)** Tumor growth curves and final tumor weights of Rv1-WT xenografts growing in intact or castrated mice, treated with SNX631 (30 mg/kg, b.i.d.) or vehicle via oral gavage. **(D)** Effects of SNX631 treatment (30 mg/kg, BID) on body weight changes for the animals in the Rv1-WT studies (C). **(E)** Representative images of H&E staining and machine-learning based coloring of the sections of endpoint tumors collected from Rv1-Luc studies (B). Coloring: proliferation zone (green), apoptotic zone (red), necrotic zone (yellow). **(F)** Quantitative analysis of the area percentage of the indicated zones in different tumor sections. **(G)** Castrated Ncr nude mice were treated with suspension vehicle or SNX631 (in suspension vehicle) at 25 mg/kg b.i.d., 50 mg/kg q.d., or 50 mg/kg b.i.d. for 39 days; tumor volumes monitored for a total of 300 days from start of treatment. Top: tumor growth curves (left) and mouse body weight changes (right). Bottom: KM plot of event-free survival (left) and spider plot of tumor growth in individual animals (right). **(H)** Castrated Ncr nude mice were treated with solution vehicle or SNX631 (in solution vehicle) at 15 mg/kg BID, 30 mg/kg QD, or 30 mg/kg BID for 63 days and then treated with control or SNX631-medicated diet (350 ppm) for up to 300 days since the start of treatment; plots are the same as for (**G**).

We have investigated the long-term effects of Mediator kinase inhibition by systemic treatment with SNX631 in Rv1-WT xenografts growing in castrated NCr nude mice, which, unlike NSG, contain NK cells that are known to be stimulated by CDK8/19i.^20,21^ In the first study (Fig. 5G), mice were treated with SNX631 over 38 days. CDK8/19i at first slowed down tumor growth relative to the control group but then tumors stopped growing and some regressed (such regression was not seen in NSG mice); the treatment had no detrimental effect on mouse body weights. After 38-day treatment, mice were monitored, and tumor sizes measured for a total of 300 days. Some tumors resumed their growth after the cessation of treatment, but others shrank (Fig. 5G). 5/30 tumors (16.7%) disappeared and have not recurred for the rest of the 300-day period, indicating the achievement of cures. In the second study (Fig. 5H), SNX631 treatment was continued for the entire 300-day period. There were no detectable adverse effects during the whole 300-day treatment period, although a few mice (1 of 11 in the vehicle group and 3 of 29 in the SNX631 group) died for treatment-unrelated reasons. In that study, complete tumor disappearance without recurrence was observed in 7 of 28 tumors (25%).

### Transcriptomic analysis of the in vivo effects of CDK8/19 inactivation in 22Rv1 xenografts

To understand why Mediator kinase inhibition or mutagenesis strongly inhibit 22Rv1 growth *in vivo* in castrated mice but have only weak effects in intact animals, we have carried out RNA-Seq analysis of tumors formed in intact and castrated NSG mice by Rv1-WT and Rv1-Luc, with and without SNX631 treatment, and by dKO-19 and dKO-19M derivatives. RNA-Seq data were analyzed separately for the human (tumor) and stromal (mouse) RNA as described.^49^ The numbers of tumor-derived DEGs obtained in different comparisons (FC >1.5, FDR <0.01 cutoffs) are shown in the volcano plots in Fig. S3A-C. Remarkably, the number of genes upregulated by Mediator kinase inhibition was much greater in tumors growing in castrated than in intact mice (8.2-fold in Rv1-WT, 4.8-fold in Rv1-Luc and 4.6-fold in dKO-19M/dKO-19); the number of downregulated genes also increased 1.6-to 2.5-fold in castrated mice (Fig. S3A-C). Hence, broader transcriptomic effects of Mediator kinase inhibition in tumors growing in castrated mice may underlie the preferential suppression of such tumors.

In contrast to similar effects of Mediator kinase inhibition on gene expression in 22Rv1 cells growing in FBS and CSS media (Fig. 3C, D), the majority of Mediator kinase affected DEGs in tumor cells growing in castrated animals were not affected in tumor cells growing in intact mice or *in vitro* (Fig. S3D). On the other hand, a high fraction of the DEGs affected by Mediator kinase inhibition in castrated mice were also affected by castration (27-56% in different 22Rv1 models) (Fig. S3E). The number of DEGs upregulated by castration was increased 1.3-to 2.0-fold under the conditions of Mediator kinase inhibition (Fig. S3A-C), suggesting that Mediator kinase activity could restrain castration-induced transcriptomic changes.

The effects of castration on MYC expression (Fig. S4A) matched its effects on tumor growth (Fig. 4A), decreasing MYC in Rv1-WT and, to a lesser extent, in dKO-19, but not in castration-resistant Rv1-Luc. Mediator kinase inhibition or mutagenesis, however, had no significant effect on MYC expression (Fig. S4A). AR expression was not affected by CDK8/19 inhibition in intact mice but was inhibited in Rv1-Luc and even more in dKO-19 tumors in castrated mice. In Rv1-WT, however, AR expression was strongly inhibited by castration alone and was not further reduced by CDK8/19i treatment, indicating that it was not a general mechanism of the CDK8/19 effect.

GSEA analysis showed that Mediator kinase inhibition or mutagenesis affected 12 of 50 hallmark pathways (Fig. 6A). Several pathways were downregulated in all three models, including those related to cell proliferation (E2F and MYC targets, G2/M checkpoint), as well as the oxidative phosphorylation pathway. The androgen response pathway was affected by castration but not by CDK8/19i treatment. Several pathways were upregulated by Mediator kinase inhibition, and most of them were also enhanced by castration (Fig. 6A). The latter pattern was especially noticeable among the transcription factor pathways affected by Mediator kinase inhibition (from C3 transcription factor targets legacy collection) (Fig. 6B). Mediator kinase inactivation in castrated mice and castration under the conditions of Mediator kinase inactivation had very similar effects on these pathways, most of which were upregulated, and some (E2F) downregulated by both treatments. The effect of castration on transcription factor pathways was greatly increased by Mediator kinase inhibition, providing a further indication that Mediator kinase activity restrained the transcriptional effects of castration.

**Figure 6.**
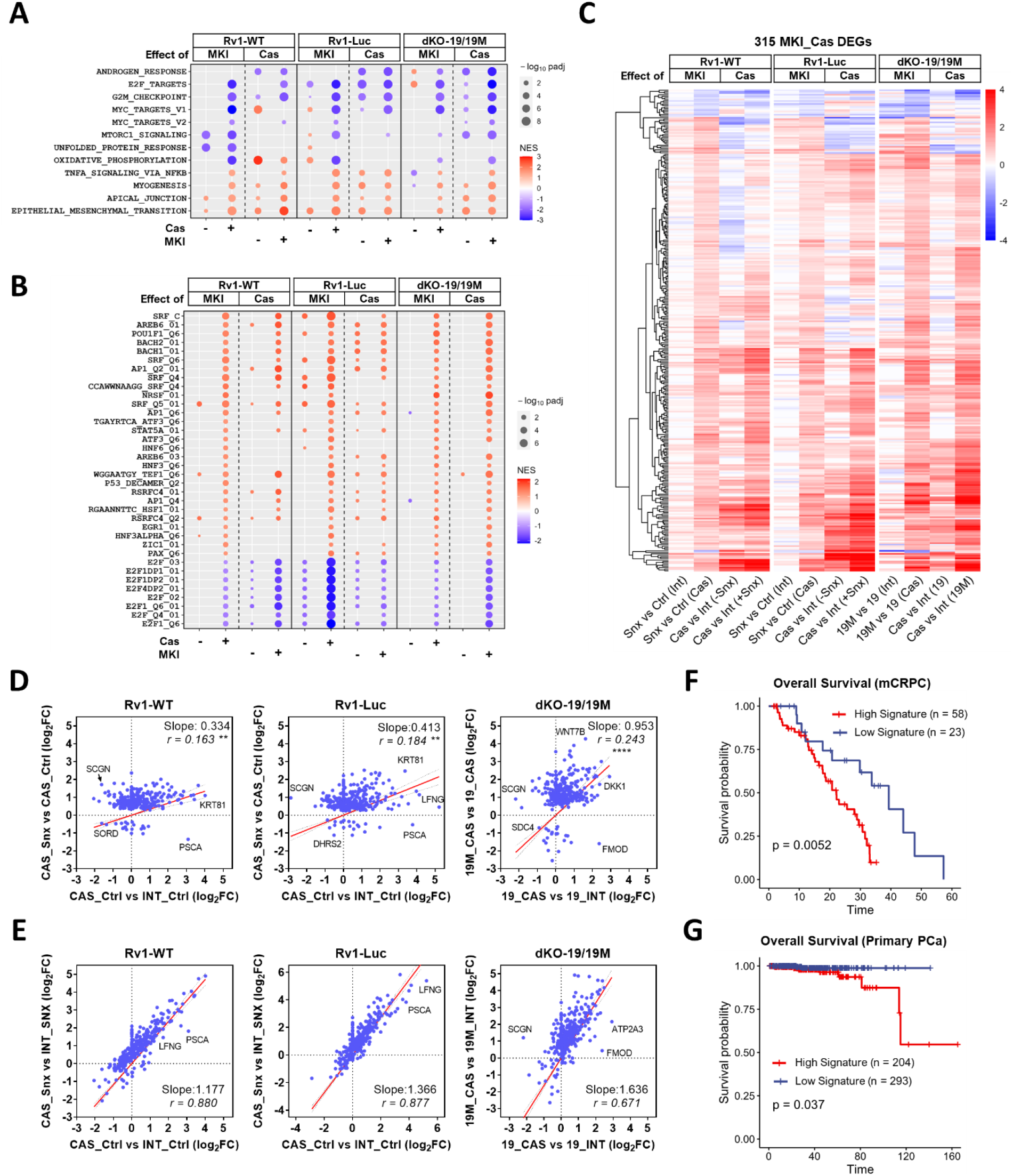
Transcriptomic analysis of the effects of Mediator kinase inhibition on tumor (human) genes in 22Rv1 xenografts. **(A)** Effects of Mediator kinase inhibition (MKI) by SNX631 treatment (Rv1-WT, Rv1-Luc) or mutagenesis (dKO-19/19M) and of castration (Cas) under the indicated conditions on the pathways affected by MKI in three 22Rv1 tumor models in castrated animals. Affected pathways were identified by GSEA of the Hallmark pathways. **(B)** Same as **(A)**, for C3 transcription factor targets signature gene sets (MSigDB Human collections). **(C)** Heatmap of 315 tumor DEGs commonly co-regulated by MKI in three 22Rv1 models in the indicated comparisons. **(D)** Correlation of the effects of MKI in castrated animals and the effects of castration on the same DEGs. (**, p<0.01; ****, p<0.0001). **(E)** Correlation of the effects of castration with and without MKI on the same DEGs. **(E,F)** Correlations of the gene signature comprised of 266 DEGs (expressed in clinical samples; out of the 315 DEG set) with overall survival in mCRPC patients **(E)** and primary PCa patients **(F)**.

To identify genes that may be involved in tumor suppression of all three 22Rv1 models in castrated mice, we have selected 315 DEGs that were co-regulated by SNX631 treatment in both Rv1-WT and Rv1-Luc tumors and were also differentially expressed between dKO-19M and dKO-19 tumors in castrated mice (Table S3). The effects of Mediator kinase inhibition or castration on the expression of these genes are shown in the heatmap in Fig. 6C. Only 20 (6.3%) of these DEGs were downregulated by Mediator kinase inhibition while the others were upregulated. The effects of Mediator kinase inhibition or mutagenesis in castrated mice and the effects of castration under the conditions of Mediator kinase inactivation on these 315 DEGs were strikingly similar (Fig. 6C). There was a significant correlation between the effects of castration and the effects of Mediator kinase inhibition in castrated mice on these DEGs (Fig. 6D). Fig. 6E shows that the effect of castration on these genes was much stronger under the conditions of Mediator kinase inhibition (as indicated by slope > 1), indicating that Mediator kinase inhibition largely enhanced the transcriptomic effects of castration in the tumor cells (although a few genes showed opposite responses to castration and Mediator kinase inhibition). The hyper-activation of castration-inducible genes by Mediator kinase inhibition resembles the hyper-induction of super-enhancer-associated genes upon Mediator kinase inhibition described in AML cells.^19^

Fig. S4B,C show the effects of different treatments, *in vitro* and *in vivo*, on the expression of selected DEGs that represent distinct patterns of response to castration and Mediator kinase inhibition. Among the genes upregulated by castration but downregulated by Mediator kinase inhibition, we note PCa biomarkers PSCA^50^ and FMOD,^51^ whereas genes such as SORD (an androgen-responsive gene^52^) and ENSG00000289695 are downregulated both by castration and by Mediator kinase inhibition (Fig. 4B). In the much larger category of genes that are induced by castration and hyper-induced when castration is combined with Mediator kinase inhibition (Fig. S4C), we note an AR-regulated WNT protein WNT7B^53^ and WNT inhibitor DKK1^54^, as well as annexins ANXA1 and ANXA3, keratins KRT15, KRT17 and KRT81, and LFNG, implicated in tumor suppression in prostate cancer.^55^ Interestingly, ETV6 and FOSL2, two of six genes identified in AML^19^ as associated with super-enhancers and hyper-activated by Mediator kinase inhibition with the resultant inhibition of cell proliferation, were also upregulated by Mediator kinase inactivation in 22Rv1 tumors in castrated mice (Fig. S4D).

We have asked whether the expression of genes regulated by Mediator kinases in 22Rv1 tumors in castrated mice could affect patient survival in mCRPC. To this end, we have generated a gene signature indicative of Mediator kinase activity under the conditions of castration, comprising 266 (out of 315) DEGs (Table S4) that were expressed (fpkm > 0.1) in mCRPC clinical samples. We then tested whether this signature correlates with overall survival (OS) in the RNA-Seq dataset of 81 mCRPC patients for whom the survival data were available.^36^ The Mediator kinase activity signature showed a strong negative correlation with OS among mCRPC patients (Fig. 6E). Interestingly, this signature also correlates negatively with OS among 497 primary PCa patients (Fig. 6G), even though the latter samples were collected prior to castration.

We have also analyzed the effects of Mediator kinase inhibition on stroma-derived (mouse) genes in the three 22Rv1 tumor models. Since the number of mouse reads from these tumors was low relative to the human reads (Fig. S5A), our statistical analysis of stromal genes was limited but yielded interesting observations. We have identified 97 stromal DEGs that were affected by SNX631 treatment of 22Rv1-WT and 22Rv1-Luc tumors in castrated mice (using p<0.05 and FC >1.5 as cutoff criteria). Surprisingly, ½ of these genes (48 DEGs) were affected not only by systemic treatment with SNX631 but also by Mediator kinase mutation in tumor cells alone, suggesting that Mediator kinase activity in tumor cells is involved in molding the tumor microenvironment. A heatmap of the effects of different treatments on these DEGs is shown in Fig. 7A, whereas Fig. 7B shows the heatmap of 49 genes that were affected by systemic treatment with SNX631 but not by Mediator kinase mutation in tumor cells. As in the case of Mediator kinase-regulated tumor genes, the effects of Mediator kinase inhibition and castration on the stromal gene sets were correlated in 2 of 3 models (Fig. 7C), and Mediator kinase inhibition increased the effects of castration on these genes (slope >1) (Fig. 7D). The expression of selected stromal genes representing different regulatory patterns is shown in Fig. S5B,C. We note that changes in some of these genes could have contributed to the inhibition of tumor growth in castrated mice, such as upregulation of Igfbp4^56^, Ccdc80^57^ and Dlk1^58^ and downregulation of Ramp3^59^ or Osm.^60^

**Figure 7.**
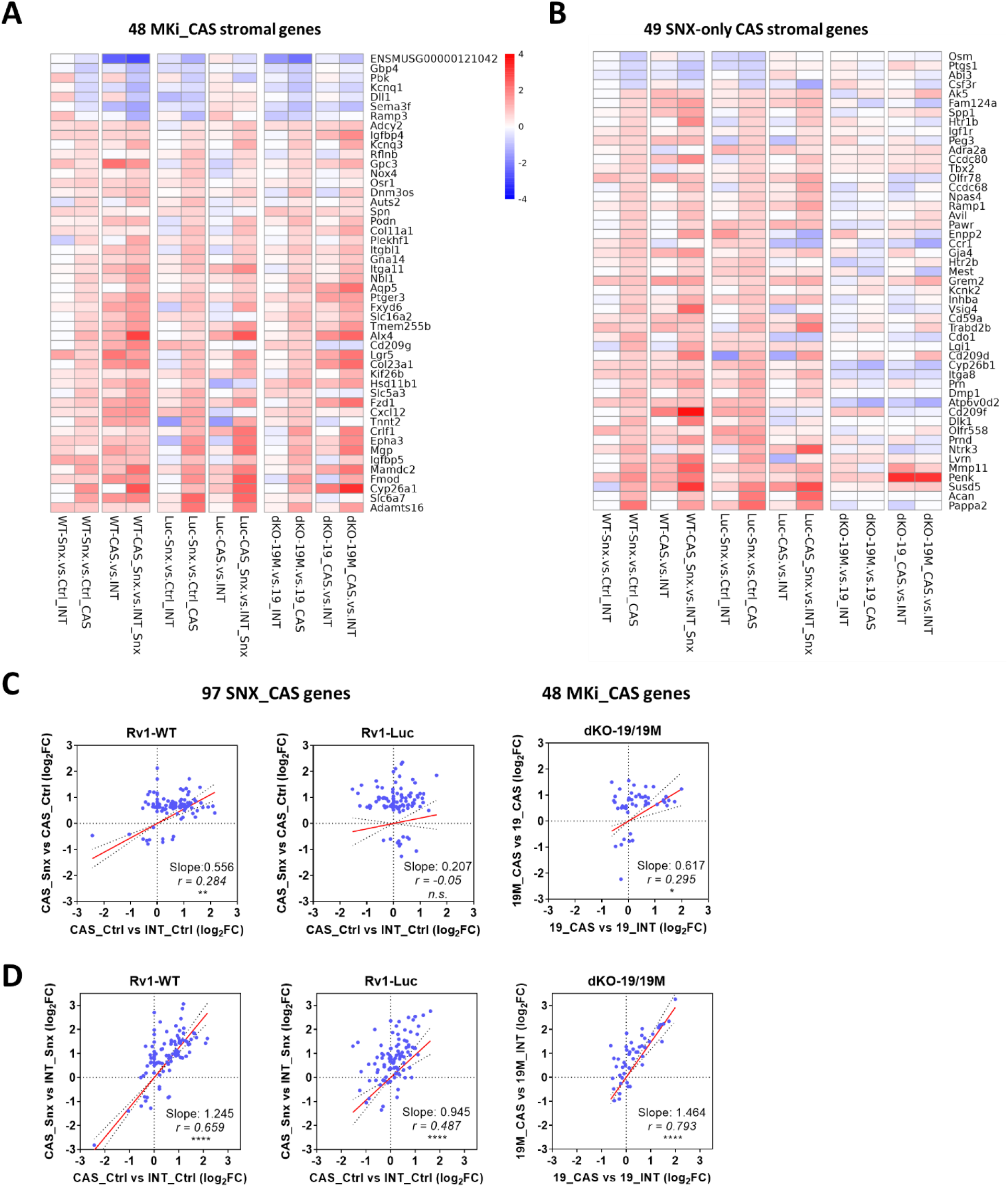
Transcriptomic analysis of the effects of Mediator kinase inhibition on stromal (mouse) genes in 22Rv1 xenograft tumors. **(A)** Heatmap of 48 stromal DEGs affected by both SNX631 treatment and Mediator kinase mutagenesis in castrated animals under the indicated conditions. **(B)** Heatmap of 49 stromal DEGs affected by SNX631 treatment but not by Mediator kinase mutagenesis under the indicated conditions. **(C)** Correlation of the effects of MKI in castrated animals and the effects of castration on stromal DEGs regulated by Mediator kinase mutagenesis and/or SNX631 treatment. (n.s., not significant; *, p<0.05; **, p<0.01). **(D)** Correlation of the effects of castration with and without MKI on stromal DEGs regulated by Mediator kinase mutagenesis and/or SNX631 treatment. (****, p<0.0001).

### Effects of CDK8/19 inhibition on patient-derived xenograft (PDX) CRPC models

We next investigated if Mediator kinase inhibition would affect the growth of three clinically relevant patient-derived xenograft (PDX) models of AR-positive CRPC. The first two, SM0310 and CG0509 were derived from adenocarcinomas of patients who failed both ADT and chemotherapy (Fig. S6A) and displayed positive AR immunostaining (Fig. S6B). SNX631 treatment inhibited tumor growth of SM0310 in both intact (Fig. 8A) and castrated (Fig. 8B) male NSG mice. Remarkably, prolonged treatment with SNX631 (67 days) stabilized the PDX growth in castrated mice once the size of xenograft reached ∼1,000 mm^3^ (Fig. 8B). CG0509 showed poor tumor take in castrated mice but its growth in intact NSG mice was inhibited after 53 days of treatment with SNX631 (Fig. 8C).

**Figure 8.**
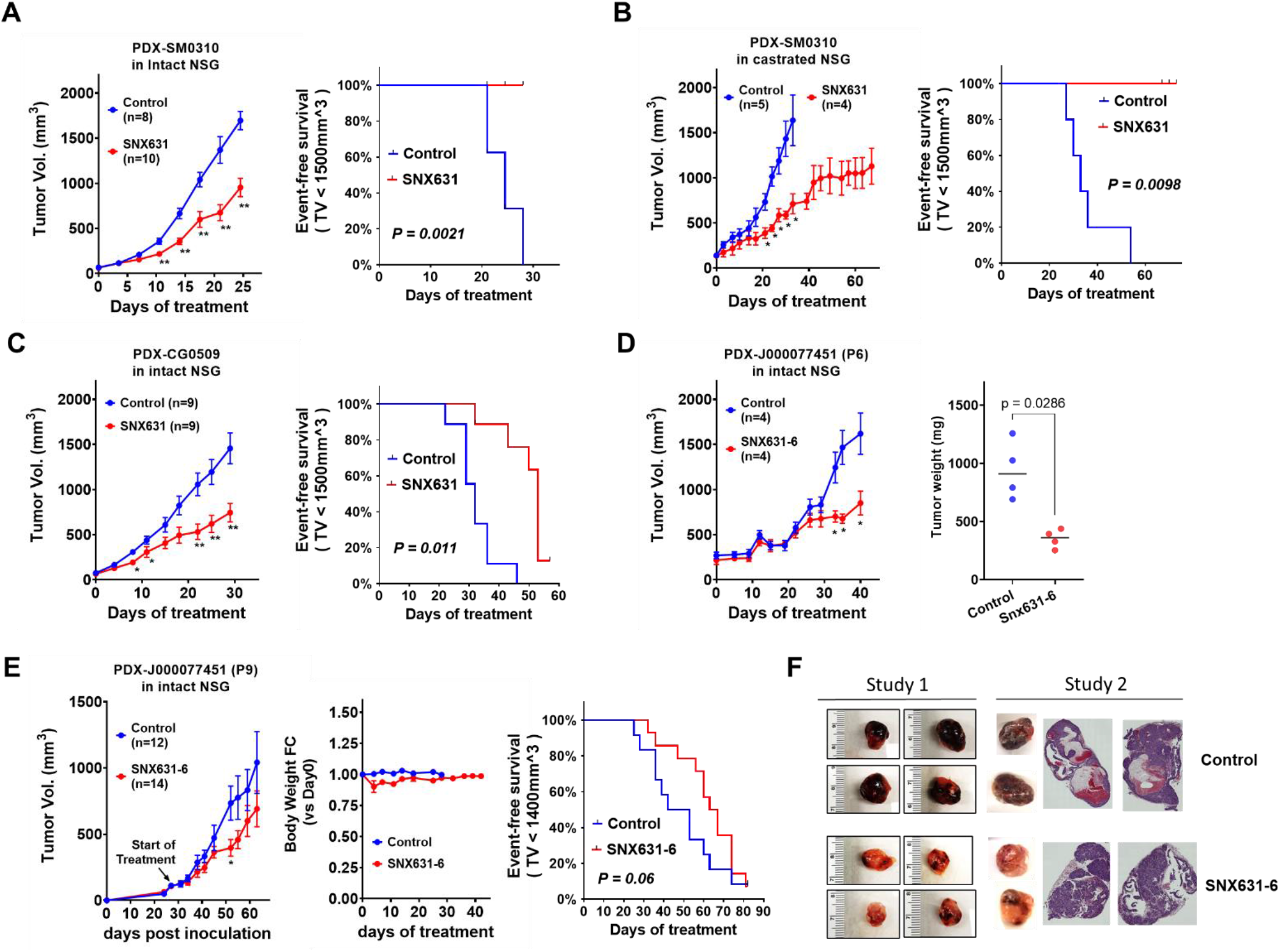
Effects of Mediator kinase inhibitors on PDX CRPC growth. **(A)** Effects of SNX631 treatment (medicated food, 500 ppm) on tumor growth curves and event-free survival of SM0310 PDX model in intact NSG mice (event defined as tumor volume exceeding 1.5 cm^3^). **(B)** Effects of SNX631 treatment on tumor growth curves and event-free survival of SM0310 PDX in castrated NSG mice. **(C)** Effects of SNX631 treatment on tumor growth curves and event-free survival of CG0509 PDX model in intact NSG mice. **(D)** Effects of SNX631-6 treatment (medicated food, 500 ppm) on tumor growth curves and final tumor weights of J000077451 PDX model in intact NSG mice (study 1). **(E)** Effects of long-term SNX631-6 treatment on tumor growth curves, mouse body weights and event-free survival of J000077451 PDX model in intact NSG mice (Study 2). **(F)** Macroscopic images and H&E staining of representative control and SNX631-6-treated J000077451 PDX tumors from **D** (study 1) and **E** (study 2).

The third PDX, J000077451 (Jackson Labs) was derived from brain metastasis of grade IV prostate adenocarcinoma. While the patient’s treatment history is unknown, the growth of this PDX is resistant to cisplatin and weakly inhibited by docetaxel (Jackson Labs website). This PDX had a low tumor take rate in castrated mice; therefore, the studies were conducted in intact male NSG mice. For Mediator kinase inhibition in this model, we used SNX631-6, an equipotent analog of SNX631. (Figure S7 shows the structure of SNX631-6 (Fig. S7A), its CDK8/19 selectivity based on kinome profiling (Fig. S7B), its potency in a CDK8/19-dependent cell-based assay (Fig. S7C) and cell-free target binding kinetics data for CDK8 and CDK19 (Fig. S7D)). In the first study, SNX631-6 treatment did not affect J000077451 tumor growth for the first 26 days of treatment but strongly inhibited the PDX growth afterwards, as also evidenced by the final tumor weights (Fig. 8D). In the second study, SNX631-6 treatment extended the median event-free survival by 37% (65 days vs 47.5 days) (Fig. 8E). The control J000077451 tumors displayed a high blood content, as indicated by a dark colored hemorrhagic phenotype while the SNX631-6 treated tumors were lighter colored and less hemorrhagic (Fig. 8F, left). H&E staining showed that the control tumors contained cavities filled with blood cells, which were not observed in SNX631-6 treated tumors (Fig. 8F, right), suggesting that the delayed tumor-suppressive effect of Mediator kinase inhibitor in this model could be due to the interference with tumor vasculature.

## DISCUSSION

The incurability of advanced PCa is largely attributed to the remarkable plasticity of CRPC cells and their propensity for cellular reprogramming that underlies such plasticity.^3^ Our findings suggest that Mediator kinase paralogs, CDK19 and CDK8, which function as broad-spectrum regulators of transcriptional reprogramming,^8^ play a key role in the ability of CRPC to adapt to the conditions of androgen deprivation *in vivo*. As a result, PCa progression and the acquisition of androgen independence render CRPC dependent on the Mediator kinase activity for its *in vivo* growth. The Mediator kinase dependence of CRPC offers a novel therapeutic opportunity for suppressing CRPC growth and even producing cures by treatment with Mediator kinase inhibitors.

The involvement of Mediator kinases in PCa progression likely reflects their inherent role in prostate tissue physiology. The prostate and the other androgen-dependent organ, the testis, have the highest CDK19 expression among normal tissues; the testis is also the highest expressor of CDK8. Our finding that androgen downregulates CDK8 while it upregulates CDK19 in androgen-responsive PCa cells offers a rare example of the regulation of Mediator kinase expression by a physiological agent. The analysis of Mediator kinase expression in clinical cancers, together with the prior reports,^26^ shows that CDK8 is downregulated and CDK19 upregulated in primary PCa. We can now interpret these findings through the effect of increased androgen signaling in PCa on the transcription of CDK8 and CDK19. (In contrast to the prostate, testicular carcinogenesis is associated with a uniquely strong downregulation of both CDK8 and CDK19. Perhaps not coincidentally, testicular carcinoma is also the most chemotherapy-curable cancer in adults.) Downregulation of CDK8 ceases, however, when PCa progresses to mCRPC and canonical androgen signaling is altered, at which stage CDK8 becomes strongly upregulated, along with a further increase in CDK19 and all the other components of the kinase module of the Mediator complex (CCNC, MED12, MED13, MED13L). Concordant upregulation of all the subunits of this module during PCa progression argues that such upregulation is not incidental to other transcriptional changes but is driven by selection for increased Mediator kinase activity.

Analysis of the transcriptomic effects of Mediator kinase inhibition on androgen-regulated transcription in androgen-responsive LNCaP cells shows that CDK8/19i both counteract the induction of the most strongly androgen-responsive genes (including KLK3 (PSA)) and enhance the effect of androgen on a subset of genes. Mediator kinase inhibition does not inhibit the mitogenic effects of androgen on LNCaP cells *in vitro* but mildly reduces LNCaP cell proliferation under androgen-deprived conditions. In concordance with this, *in vitro* proliferation of an LNCaP derivative C4-2, which was adapted to growth in androgen-depleted media, was inhibited by CDK8/19i only when AR signaling was suppressed, in agreement with a prior study in LNCaP cells.^31^ CDK8/19i treatment also moderately inhibited the growth of C4-2 *in vivo* in intact male mice but we could not use this model to investigate the effect of CDK8/19 inhibition under the androgen deprivation conditions *in vivo* due to the low C4-2 tumor take rate in castrated mice.

Instead, we have focused on 22Rv1, a “classical” CRPC model, to analyze the role of CDK8 and CDK19 in tumor growth with and without androgen deprivation, through both genetic modifications and pharmacological inhibition of CDK8/19. Analysis of two 22Rv1 cell lines with wild-type CDK8/19 that differ in their response to ADT and of a series of 22Rv1 derivatives expressing wild-type or kinase-inactive mutants of CDK8 or CDK19 showed that the *in vitro* transcriptomic effects of CDK8 and CDK19 were qualitatively similar, in agreement with the previous study.^8^ While 22Rv1 proliferation *in vitro* was moderately inhibited by Mediator kinase inhibition, 22Rv1 tumor growth *in vivo* was dramatically suppressed by Mediator kinase inhibition or mutagenesis in castrated mice but Mediator kinase-deficient tumors still grew in intact male animals. Furthermore, genetic inactivation of CDK8/19 rendered tumors formed by these CRPC cells responsive to a conventional ADT agent (degarelix). Hence, Mediator kinase activity is required for CRPC growth *in vivo* under the conditions of androgen deprivation.

The mechanism of the selective effect of Mediator kinase mutagenesis or systemic inhibition on *in vivo* growth of 22Rv1 CRPC in castrated mice was investigated by RNA-Seq analysis. This analysis showed that the transcriptomic effects of Mediator kinase inhibition were very different between the tumor cells growing in castrated or in intact male mice (or in cells growing in culture). In particular, Mediator kinase inhibition or mutagenesis have affected (mostly upregulated) a much greater number of genes when tumors grew in castrated mice. More than one-third of the tumor genes regulated by Mediator kinases in castrated animals were affected by castration alone, and the number of castration-affected genes was increased when Mediator kinases were inhibited. A few of these genes (such as PSCA, a positive regulator of PCa growth and metastasis^61^) were upregulated by castration but downregulated by Mediator kinase inhibition, in agreement with the previously reported inhibition of stress-induced transcription by CDK8/19i.^9^ The largest group of the CDK8/19-regulated genes, however, was induced by castration and upregulated even more by Mediator kinase inhibition or mutagenesis. In addition, Mediator kinase inactivation strongly increased the number of transcription factor pathways that were affected (mostly upregulated) by castration. These results suggest that Mediator kinase activity restrains castration-induced transcriptional reprogramming, as has been previously shown for its effect on chemically induced cell-fate reprogramming.^12^

Potentiation of castration-induced gene expression by Mediator kinase inhibition closely resembles the hyper-induction of super-enhancer-associated genes upon CDK8/19 inhibition in AML cells, where the resulting unbalanced expression of such genes inhibited cell proliferation.^19^ Interestingly, two out of six CDK8/19i-hyper-activated genes in AML cells, upregulation or downregulation of which inhibited AML cell growth,^19^ were also upregulated by Mediator kinase inhibition in 22Rv1 tumors. It remains to be determined whether the genes induced by castration and Mediator kinase inhibition in CRPC are also associated with super-enhancers. This seems possible, since super-enhancers characterized in PCa are enriched in the Mediator complex,^62^ and the levels of this complex are increased by Mediator kinase inhibition.^8^ Tumor suppression by either downregulation or upregulation of key transcriptional signals is well established in PCa, where both androgen deprivation and supra-physiological doses of testosterone (SupraT) suppress the growth of cancers with canonical AR signaling, the basis of bipolar androgen therapy.^63^ While CRPC with altered AR signaling (such as 22Rv1) are not sensitive to SupraT, hyper-induction of castration-responsive genes by Mediator kinase inhibition resembles the effects of SupraT in producing a strong tumor-suppressive effect.

In addition to the transcriptomic effects on tumor cells, we found that Mediator kinase inhibition affects stromal gene expression in a way that could also have contributed to tumor suppression. Several stromal genes upregulated by Mediator kinase inhibition were shown to have tumor-suppressive effects in the stroma (Igfbp4^56^, Ccdc80^57^, Dlk1^58^) and some downregulated genes were reported to have tumor-supporting activities (Ramp3^59^, Osm^60^). Importantly, many of the stromal genes were affected not only by systemic treatment with a CDK8/19 inhibitor but also by Mediator kinase mutagenesis in tumor cells alone, suggesting that CDK8/19 activity in the tumor may mold the tumor microenvironment.

The role of the tumor environment was also suggested by two long-term (300-day) studies, where 22Rv1 xenografts in castrated nude mice were treated with SNX631 either for the first 38-days or for the entire 300-day period (such an exceptional length of continuous drug treatment was made possible by the lack of toxicity of selective Mediator kinase inhibitors). Long-term treatment or long-term follow-up after a shorter treatment period revealed not only tumor growth inhibition but also regression, with 17-25% of tumors disappearing and not recurring till the end of the 300-day period, indicating the achievement of cures. To the best of our knowledge, there are no other reports of cures achieved in 22Rv1 or other CRPC models. Importantly, tumor regression and cures were observed in NCr nude mice that lack mature T-cells but still have B cells and robust NK cell responses. On the other hand, we have not seen tumor regression in NSG mice, which are B, T and NK cell deficient with impaired innate immunity. Since NK cell activity is known to be stimulated by CDK8 knockdown or CDK8/19i treatment,^20,21^ it is possible that NK cells could be involved in tumor regression upon systemic CDK8/19i treatment.

Mediator kinase inhibitors also suppressed tumor growth in three AR-positive PDX models derived from PCa patients, at least two of whom had failed both ADT and chemotherapy. Interestingly, the effect of CDK8/19 inhibition in one of these models was associated with the suppression of intratumoral blood supply, an apparent stromal effect. In a related example, positive regulation of angiogenesis by CDK8 was previously suggested for pancreatic cancer.^64^ PDX suppression was observed in intact mice, in contrast to the 22Rv1 model but in concordance with the results obtained with C4-2 cells. Hence, the requirement of castration for the effect of CDK8/19i on CRPC is not absolute but depends on the specific tumor. From the clinical standpoint, however, the responses seen under the conditions of androgen deprivation should be especially relevant, since most CRPC patients would have undergone chemical castration. Remarkably, the gene expression signature reflecting Mediator kinase activity in 22Rv1 tumors showed a strong association with overall survival in mCRPC patients. Thus our results warrant the exploration of Mediator kinase inhibitors as a new class of drugs for the treatment of CRPC, which is resistant to currently available therapies.

## STAR★METHODS

### KEY RESOURCES TABLE

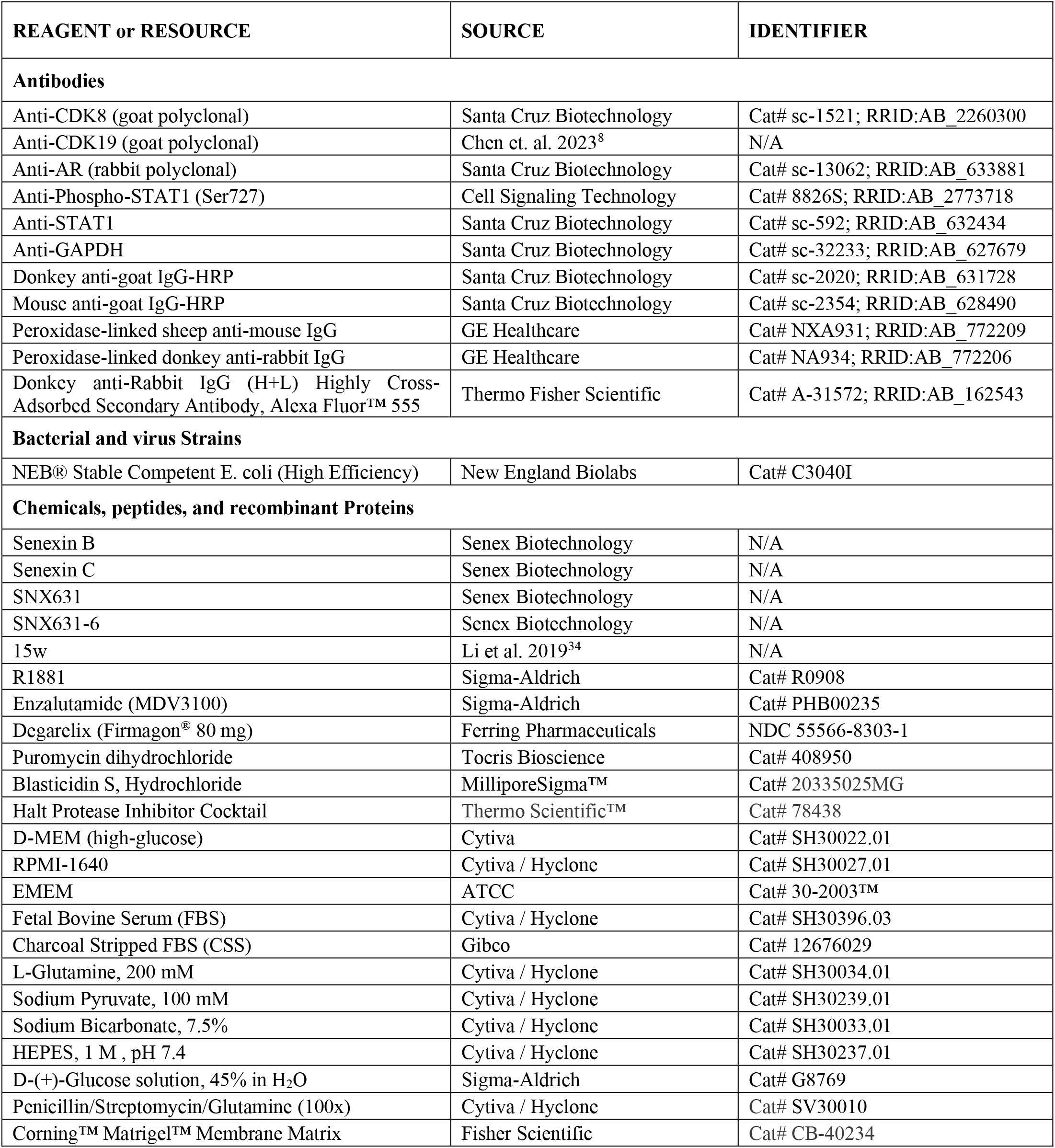
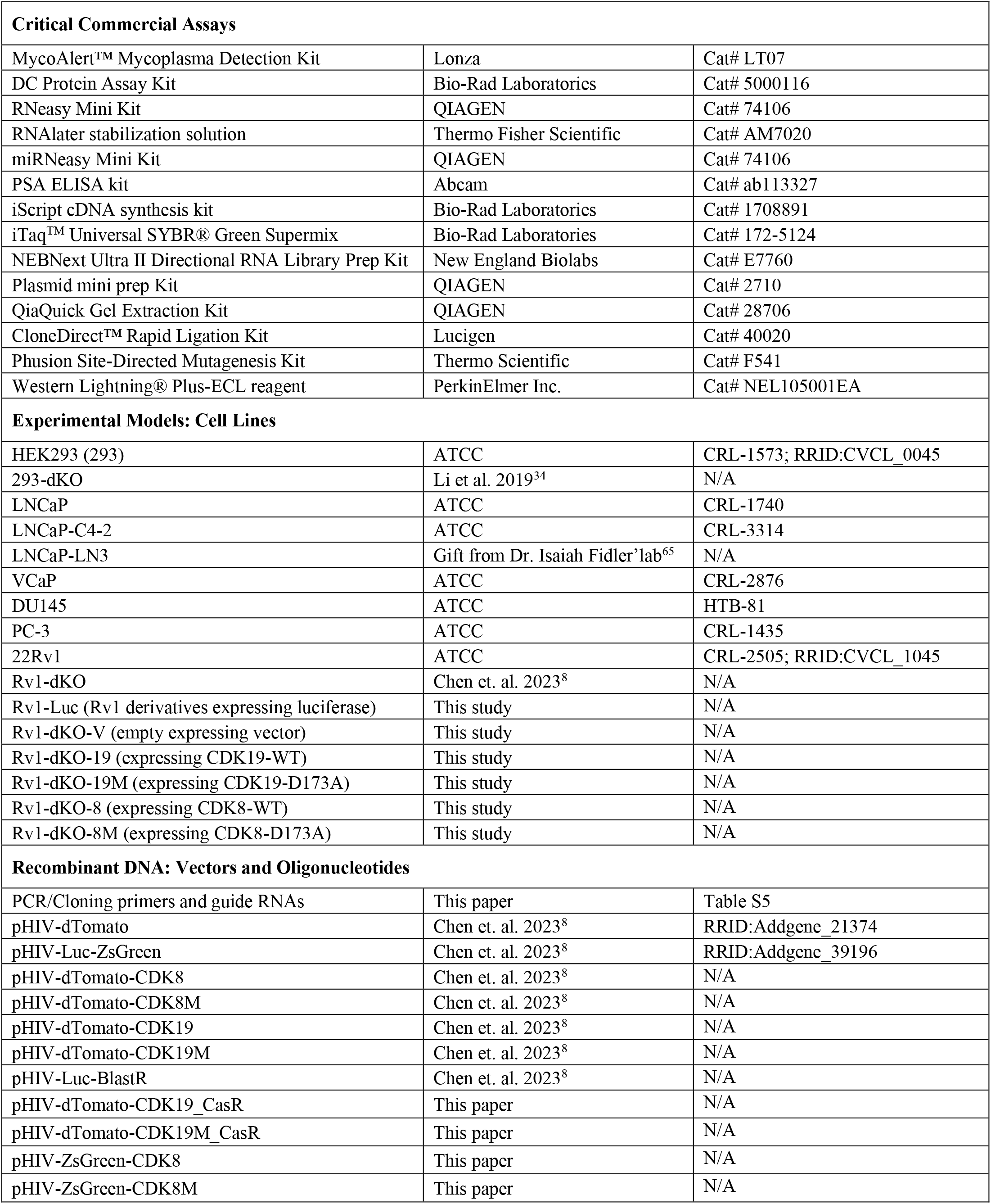
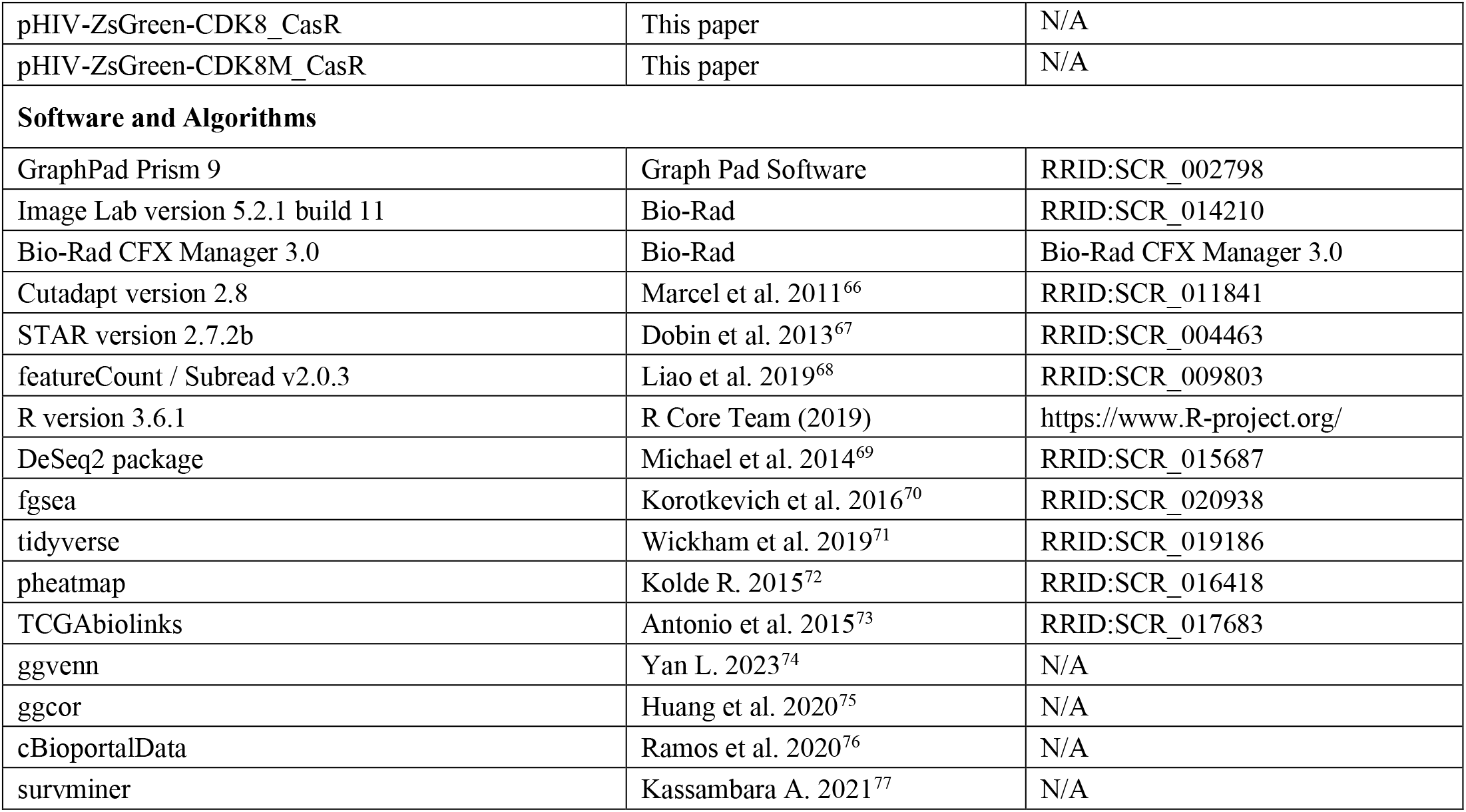

## RESOURCE AVAILABILITY

### Lead contact

Further information and requests for resources and reagents should be directed to and will be fulfilled by the Lead Contacts, Mengqian Chen or Igor Roninson.

### Materials availability

Senexin B, Senexin C, SNX631 and SNX631-6 can be obtained from Senex Biotechnology via an MTA.

Materials generated in this study are available upon request to the lead contact.

### Data and code availability

The RNA-Seq data are available in GEO database through accession number GSE240167, GSE240369 and GSE240370. Any additional information required for reanalyzing the data is available from the lead contact upon request.

## EXPERIMENTAL MODEL AND SUBJECT DETAILS

### Mice

The efficacy study in C4-2 xenograft model was performed in male NCG mice (NOD-Prkdc^em26Cd52^ /Il2rg^em26Cd^^22^/NjuCrl, Nanjing Galaxy Biopharmaceutical Co., Ltd) by Crown Bioscience (Beijing) Inc. The efficacy study in PDX-SM0310 and PDX-CG0509 models was performed in male NSG mice (NOD.Cg-Prkdc^scid^/Il2rg^tm1Wjl^/SzJ, The Jackson laboratory) at Medical University of South Carolina. All other *in vivo* animal studies were performed in male NSG or NCr nude (CrTac:NCr-Foxn1^nu^, Taconic) mice at University of South Carolina. All the experimental animal procedures were performed according to the guidelines approved by the Institutional Animal Care and Use Committee (IACUC) at the study sites. All mice were maintained under pathogen-free conditions and random assignment was employed to distribute the animals across different experimental groups in each study.

### Cell Lines

HEK293 (293), 293 derivative with CDK8/19 double-knockout (293-dKO), VCaP, LNCaP, LNCaP-C4-2, LNCaP-LN3, DU145, PC-3, 22Rv1 (Rv1-WT), and 22Rv1 derivative with CDK8/19 double-knockout (Rv1-dKO) were acquired from ATCC and other sources listed in the KEY RESOURCES TABLE. 22Rv1 were transduced with pHIV-Luc-BlastR to generate the derivative Rv1-Luc. Rv1-dKO cells were reconstituted with CDK19-wild-type or CDK19-D173A kinase-inactive mutant expression constructs or insert-free vector to generate derivatives Rv1-dKO-V, Rv1-dKO-19 and Rv1-dKO-19M. Rv1-dKO-V cells were further reconstituted with CDK8-wild-type or CDK8-D173A kinase-inactive mutant expression constructs to generate derivatives Rv1-dKO-8 and Rv1-dKO-8M. More details can be found in the METHOD DETAILS section.

### Patient-derived xenograft (PDX) models

PDX-SM0310 and PDX-CG0509 models were established at the University of California, Irvine. The PDX J000077451 model was acquired from the Jackson Laboratory (https://tumor.informatics.jax.org/mtbwi/pdxDetails.do? modelID=J000077451).

## METHOD DETAILS

### Cell culture

LNCaP and its derivatives C4-2 and LN3, as well as 22Rv1 and its derivatives, were cultured in RPMI-1640 media supplemented with 2 mM L-glutamine, 1 mM sodium pyruvate, 1.5 g/L sodium bicarbonate, 10 mM HEPES, 0.45% D-glucose, and either 10% fetal bovine serum (FBS) or charcoal stripped FBS (CSS), plus 1% penicillin-streptomycin (P/S). This combination is referred to as PCa FBS media when using FBS, and PCa CSS media when using CSS. HEK293 and its derivatives, and VCaP cells were cultured in DMEM media supplemented with 10% FBS and 1% P/S. DU145 cells were cultured in EMEM media supplemented with 10% FBS and 1% P/S. All the cells were incubated at 37°C with 5% CO_2_. PC-3 cells were cultured in F-12K media supplemented with 10% FBS and 1% P/S. Regular testing for mycoplasma contamination was conducted every 3-4 weeks using the MycoAlert PLUS mycoplasma detection kit (Lonza).

### Generation of Rv1-Luc and Rv1-dKO CDK8/19 re-expression derivatives

22Rv1 cells were transduced with pHIV-Luc-BlastR lentiviral construct and selected with Blasticidin (5 μg/mL) for 3 weeks to obtain 22Rv1-Luc cells. Rv1-dKO cells, previously generated using lentiCRISPR-Puro-sgCDK8 and lentiCRISPR-Blast-sgCDK19 constructs,^8^ were transduced with pHIV-dTomato (control vector), pHIV-dTomato-CDK19_CasR and pHIV-dTomato-CDK19M_CasR lentiviral constructs and sorted for fluorescence-positive cells (two rounds) using FACS Aria III (BD Biosciences) at the Microscopy and Flow Cytometry Core (MFCC) of the CTT to obtain Rv1-dKO-V, Rv1-dKO-19 and Rv1-dKO-19M cells. Rv1-dKO-V cells were transduced with pHIV-ZsGreen-CDK8_CasR and pHIV-pHIV-ZsGreen-CDK8M_CasR lentiviral constructs, sorted by flow cytometry and passaged for clonal expansion. The clones with validated re-expression of CDK8 or CDK8M were pooled together and designated as Rv1-dKO-8 and Rv1-dKO-8M cells. The lentiviral constructs pHIV-dTomato-CDK19_CasR and pHIV-dTomato-CDK19M_CasR that contain the coding sequences of human CDK19-wild-type or CDK19-D173A kinase-inactive mutant with mutations introduced in the sgRNA-targeting sequence and proximal PAM sequences (blocking sgRNA targeting but not changing the coded protein sequences) were generated using the Phusion Site-Directed Mutagenesis Kit (Thermo Scientific) with the hCDK19-CasR-F/R primers (Table S5) and pHIV-dTomato-CDK19/CDK19M as cloning template. To create lentiviral constructs pHIV-ZsGreen-CDK8_CasR and pHIV-pHIV-ZsGreen-CDK8M_CasR, pHIV-ZsGreen-CDK8 and pHIV-ZsGreen-CDK8M were first generated by subcloning the coding sequences of human CDK8-wild-type or CDK8-D173A kinase-inactive mutant from constructs pHIV-dTomato-CDK8/CDK9M and then used as template for mutagenesis cloning with hCDK8-CasR-F/R primers (Table S5). All the lentiviruses were produced by the functional genomics core (FGC) of the University of South Carolina’s Center for Targeted Therapeutics (CTT) with a standard protocol as described previously.^8^

### In vitro cell growth assays

LNCaP-C4-2 and 22Rv1 derivatives were plated in P100 plates in PCa FBS or CSS media. After overnight attachment, the cells were treated with vehicle control (0.1% DMSO) or tested compounds at indicated concentrations. For long-duration experiments, the culture media, inclusive of the drugs, were refreshed every 3 days. At the end of the incubation, the cells were washed with 1X PBS, trypsinized, and harvested. Cell counts were determined using the TC20 Automated Cell Counter (Bio-Rad) using Dual-Chamber Cell Counting Slides (Bio-Rad) and Trypan Blue Dye (Bio-Rad). The effects of compounds at different concentrations on LNCaP cell growth were measured by Sulforhodamine B (SRB) assay in 96-well plates after 6 days of treatment.

### PSA ELISA assay

For the *in vitro* assays, cells were seeded into 12-well plates and incubated overnight to allow for adhesion and recovery. Cells were then exposed to different concentrations of CDK8/19 inhibitors for 72 hours. The culture media were harvested for analysis in parallel. For *in vivo* studies, mice bearing xenograft tumors were treated either with a vehicle control or CDK8/19 inhibitors and sera were collected post-treatment. PSA levels were quantified both in the conditioned media and in the mouse sera using the PSA ELISA kit (Abcam), according to the manufacturer’s instructions.

### Whole cell extracts and Western blot analysis

Control and treated cells grown in P100 plates were washed twice with cold PBS and then lysed in RIPA lysis buffer (50 mM Tris-HCl, pH 8.0; 150 mM NaCl; 5 mM EDTA; 0.5 mM EGTA; 1% Igepal CA-630 (NP-40); 0.1% SDS; 0.5% Na deoxycholate) supplemented with 1x protease/phosphatase inhibitor cocktail (Thermo Scientific #78438), 2 mM Na3VO4 and 10 mM NaF. Ice-cold lysates were briefly sonicated (5-10 seconds each time) 3 times to solubilize chromatin proteins followed by centrifugation (14,000 g, 20 min). Protein concentrations were determined using the DC protein assay (Bio-Rad). Lysate samples with the same amount of total protein (40-50 μg) were mixed with 4x Laemmli Sample Buffer (Bio-Rad #161-0737, supplemented with 2-mercaptoethanol) and electrophoresed in 4-12% Express-Plus PAGE gels in Tris-MOPS (SDS) running buffer (GenScript #M00138). Proteins were transferred onto PVDF membranes, which were then blocked with 5% non-fat milk and incubated subsequently with the primary and the secondary antibodies. Protein bands were visualized using Western Lighting Plus ECL detection reagent (Perkin Elmer, Waltham, MA, USA) and imaged with the ChemiDoc Touch^TM^ system (Bio-Rad). Image processing and densitometry analysis were conducted using ImageLab software (Bio-Rad).

### SNX631 treatment of C4-2 xenografts in NCG mice

This study was performed by Crown Biosciences (Beijing) Inc. C4-2 tumor cells (5×10^6^ cells in 0.1 mL 50% Matrigel per animal) were inoculated s.c. at the right front flank into male NCG mice (8-9 weeks old, 22-31 grams). On day 10, 20 animals were randomly allocated to two study groups. The mean tumor size at randomization was approximately 130 mm^3^. Randomization was performed based on “Matched distribution” randomization method (Study Director TM software, version 3.1.399.19). The animals in the control group were dosed with vehicle (0.05% Carboxymethyl cellulose solution, 10 mL/kg, p.o., q.d.) and the animals in the treatment group were dosed with SNX631 (a 2.5 mg/mL suspension formulation in the vehicle, 10 mL/kg, p.o., b.i.d.) at 50 mg/kg/day. Animals were continuously treated for 11 days, and the xenograft tumors were excised and weighed at the end of study. Endpoint serum samples were collected for PSA ELISA analysis.

### Tumor growth of 22Rv1 derivatives in intact and castrated NSG mice

22Rv1 and derivative cells were inoculated s.c. (2×10^6^ cells in 0.1 mL 50% Matrigel per animal) in the right flank of intact or castrated male NSG mice (9-11 weeks old). Castration was performed by either surgical orchiectomy (for Rv1-WT and Rv1-dKO studies) or Degarelix treatment (administered s.c. at 10 mg/kg monthly, for Rv1-Luc, dKO-V, dKO-8, dKO-8M, dKO-19 and dKO-19M studies), and tumor inoculation was done 10-14 days post castration. Tumor dimensions and body weights of the mice were recorded twice a week. Upon reaching the humane endpoint, tumors were collected and stored for further studies, including RNA-Seq (stabilized in RNA-later stabilization solution), IHC (fixed in 10% neutralized formalin for 24-48 hours, then stored in 75% ethanol), and protein analysis (snap frozen and stored at −80°C).

### Degarelix treatment of 22Rv1 derivative tumors in NSG mice

Rv1-Luc and 22Rv1-dKO derivative cells were inoculated s.c. (2×10^6^ cells in 0.1 mL 50% Matrigel per animal) in the right flank of male NSG mice (9-11 weeks old). Once the tumor volumes reached ∼150mm^3^, animals were randomly assigned to the control or treatment groups and treated with PBS or degarelix (administered s.c., 10 mg/kg once a month). The tumor sizes and body weights were measured 2-3 times per week till tumors reached the humane endpoints. Effects of degarelix treatment were evaluated using Kaplan-Meier analysis, wherein a tumor size over 1500 mm^3^ was considered as a survival endpoint.

### SNX631 treatment of 22Rv1 xenografts in intact and castrated NSG mice

Rv1-WT and Rv1-Luc cells were inoculated were inoculated s.c. (2×10^6^ cells in 0.1 mL 50% Matrigel per animal) in the right flank of intact or castrated male NSG mice (8-10 weeks old). Castration was performed by either surgical orchiectomy (for Rv1-WT study) or Degarelix treatment (administered s.c. at 10 mg/kg monthly, for Rv1-Luc study), and tumor inoculation was done 10-14 days post castration. Animals were randomly allocated to two study groups when the mean tumor size reached 100-200 mm^3^. For Rv1-WT study, the treatment group received SNX631-medicated diet (500 ppm, producing 30-60 mg/kg/day dosage on average) and the control group received the control diet. For Rv1-Luc study, the animals in the control group were dosed with vehicle (70% PEG-400/30% Propylene Glycol, 5 mL/kg, p.o., b.i.d.) and the treatment group was dosed with SNX631 (6 mg/mL solution in the vehicle, 5 mL/kg, p.o., b.i.d.) at 60 mg/kg/day. Tumor dimensions and mouse body weights were measured 2-3 times a week. At the endpoints, tumors were collected and stored for further studies, including RNA-Seq (stabilized in RNA-later stabilization solution), IHC (fixed in the 10% neutralized formalin for 24-48 hours, then stored in 75% ethanol), and protein analysis (snap frozen and stored at −80°C).

### Long-term SNX631 treatment of 22Rv1 xenografts in castrated nude mice

Rv1-WT cells were inoculated s.c. (2×10^6^ cells in 0.1 mL 50% Matrigel per animal) in the right flank of castrated male NCr nude mice (8-10 weeks old). Animals were randomly allocated to 4 study groups (one control group and three treatment groups at different dosages) when the mean tumor size reached 100-200 mm^3^. In the first long-term study, the control group animals were dosed with vehicle (0.05% Carboxymethyl cellulose solution, 10 mL/kg, p.o., q.d.) and the animals in the treatment groups were dosed with SNX631 (suspension in the vehicle at 2.5 or 5 mg/mL, 10 mL/kg, p.o., q.d. or b.i.d.) at 50 or 100 mg/kg/day. Animals were treated for 38 days and left untreated for the rest of the study period. In the second long-term study, the control group animals were dosed with vehicle (70% PEG-400 / 30% Propylene Glycol, 5 mL/kg, p.o., b.i.d.) and the animals in the treatment groups were dosed with SNX631 (solution in the vehicle at 3 or 6 mg/mL, 5 mL/kg, p.o., q.d. or b.i.d.) at 30 or 60 mg/kg/day. After 29 days of treatment, the control group animals were switched to q.d. dosing regimen and animals of all the treatment groups were switched to 30 mg/kg/day q.d. dosing regimen for another 34 days and then were put on control or SNX631-mediated chow (at 350 ppm), respectively, for the remaining period of the study. The tumor sizes and body weights were measured 2-3 times per week till tumors reached the humane endpoints. The effects of long-term SNX631 treatment were assessed using Kaplan-Meier analysis.

### Efficacy studies of patient-derived xenograft (PDX) models

For efficacy studies of PDX-SM0310 and PDX-CG0509 models, PDX tissues are dissociated into single-cell suspensions by combining mechanical dissociation with enzymatic degradation of the extracellular matrix. The tumor tissue was enzymatically digested using the Tumor Dissociation Kit (Miltenyi Biotec) and the GentleMACS™ Octo Dissociator with Heater was used for the mechanical dissociation steps. After dissociation, the sample was passed through a 70 µm strainer and washed with 20 mL of RPMI 1640 medium. The resulting single cell suspension was inoculated s.c. (1×10^6^ cells in 0.1 mL 50% Matrigel per animal) in the right flank of male NSG mice. For PDX-SM0310 studies, animals were chemically castrated with degarelix (administered s.c. at 10 mg/kg) two weeks post tumor inoculation. Animals were randomly allocated to two study groups when the mean tumor size reached 50-100 mm^3^. The treatment group received SNX631-medicated diet (500 ppm) and the control group received control diet. The tumor sizes and body weights were measured 2-3 times per week till tumors reached the humane endpoints. The effects of long-term SNX631 treatment were assessed using Kaplan-Meier analysis.

For efficacy studies of PDX-J000077451 model, male NSG mice (8-10 weeks old) were inoculated by taking a 10 mm^3^ tumor piece excised from freshly collected xenograft tissue and implanting it subcutaneously in the right flank. Once tumors reached 150 mm^3^, mice were randomized into two study groups. The treatment group received SNX631-6 in medicated diet (500 ppm, 30-60 mg/kg/day dosage on average) and the control group received control diet. Tumor dimensions and mouse body weights were measured twice a week. On the final day of the study, tumors were excised, weighed, and imaged, then stored for further studies.

### RNA extraction from cells and tumor tissues

LNCaP cells were seeded into 12-well plates at the density of 3 X 10^5^/well and cultured in PCa CSS media for 72 hours. In study 1, cells were treated (in triplicate) with different concentrations of R1881 (0.1, 1, 10 nM) for 24 hours. In study 2, cells were treated with DMSO (0.1%), Senexin B (1 μM), R1881 (0.1 nM), or a combination of Senexin B and R1881 for 24 hours. In study 3, the cells were treated with DMSO (0.1%), SNX631 (500 nM), R1881 (1 nM), or a combination of SNX631 and R1881 for 72 hours. The effects of CDK8/19 inhibition on gene expression in 22Rv1 derivatives were evaluated as follows: (1) Under androgen-supplemented conditions, cells were seeded into 12-well plates at the density of 2 X 10^5^/well in PCa FBS media, incubated overnight, and then treated with 0.1% DMSO (vehicle control) or 500 nM SNX631 in FBS media for 3 days. (2) Under androgen-deprived conditions, cells were first sub-cultured in PCa CSS media for one passage (3-4 days) and then seeded into 12-well plates at the density of 3×10^5^ /well in PCa CSS media, incubated overnight, and then treated with 0.1% DMSO (vehicle control) or 500 nM SNX631 in CSS media for 3 days. RNA was extracted from cell culture using the RNeasy Mini Kit (Qiagen) according to the manufacturer’s protocol. For tumor tissue RNA extraction, tumor xenografts were dissected from euthanized animals using sterilized surgical tools. Pieces of non-necrotic tumor tissues ∼100mm^3^ were excised at the margin near midline section and submerged in 500 μL RNA-Later stabilization solution (Thermo Fisher Scientific) at room temperature. After stabilization, RNA samples were extracted from tumor tissues using the mRNeasy Mini Kit (QIAGEN) according to the manufacturer’s protocol.

### RT-qPCR (reverse transcriptase qPCR)

RNA samples, 500 ng each, were used for cDNA synthesis utilizing the iScript cDNA Synthesis Kit (Bio-Rad). Subsequently, the expression levels of target genes were quantified through iTaq Universal SYBR Green Supermix, using CFX384 Real-Time PCR Detection System (Bio-Rad). The specific primers for qPCR are listed in Supplemental Table S5. Data derived from real-time PCR were processed via the Bio-Rad CFX Manager software, to extract Ct values from the qPCR reactions. The relative expression levels of specific genes were computed using the formula: Relative Expression= 2^(Ct__Reference_-Ct__Target_ gene), where RPL13A served as the reference gene.

### RNA-seq analysis

The RNA-Seq library preparation, next-generation sequencing (NGS), and post-processing of raw data were performed by the Functional Genomics Core (FGC) of the Center for Targeted Therapeutics (CTT). RNA-Seq libraries were generated with the NEBNext Ultra II Directional RNA Library Prep Kit and sequenced on HiSeq 3000/4000 (at Genewiz, Inc., South Plainfield, NJ) or Illumina NovaSeq (at MedGenome, Inc., Foster City, CA) platforms, utilizing paired end sequencing. Demultiplexed raw reads data was initially processed with the Cutadapt software (v 2.8) to trim off the adaptor sequences and filter out the reads that possessed short insert sequences. Processed reads were then aligned to a hybrid reference genome consisting of both the human GRCh38.p13 primary assembly genome and the mouse GRCm39 primary assembly genome, utilizing STAR (v2.7.2). The alignment output BAM files were further processed by the featureCounts software from the Subreads package, with the hybrid annotation file built from the gencode.v41.annotation.gtf and gencode.vM30.annotation.gtf annotation files downloaded from GENCODE website, to generate the counts matrix data. Differential expression (DE) analysis was performed in R using the DESeq2 package. Within the DESeq2 pipeline, the normalized counts data were fit to a negative binomial distribution model using a generalized linear model (GLM) framework, and the Benjamini-Hochberg procedure was used to control the false discovery rate (FDR) for multiple testing. The log2 fold-change (logFC) was calculated by computing the base-2 logarithm of the ratio of the mean normalized expression value (transcripts per million, TPM) for cell culture samples, or the median normalized expression value (TPM) for xenograft tissues, for each group. A small pseudocount was added to the expression values of all genes before calculating logFC to avoiding division by zero. The logFC and FDR values calculated from DESeq2 pipelines were utilized to select differentially expressed genes (DEGs). To select high-confidence DEGs from multiple RNA-Seq experiments, the following criteria were applied: (1) FDR < 0.01 in all experiments; (2) average log2FC from multiple experiments > log2(1.5). The gene set enrichment analysis (GSEA) for different comparisons was conducted using the fgsea package with the specific gene sets downloaded from the Human Molecular Signatures Database (MSigDB). All raw and processed RNA-Seq data have been uploaded to GEO (see data availability section). Detailed information about individual RNA-Seq samples (sample title and description, GEO accession number) is listed in Table S6.

### Bioinformatics analysis of clinical prostate cancer samples

For the tumor-normal comparison, RNA-seq data from the TCGA and Genotype-Tissue Expression (GTEx) portals were used. The data processing and analysis features were executed in the R statistical environment. The read counts were DESeq2 normalized, followed by a second scaling normalization. The biomaRt and AnnotationDbi R packages were then used to select and annotate appropriate genes in each of the datasets. A comparison between normal and tumor samples was performed using the Mann–Whitney U test. The statistical significance cutoff was set at p <0.01.

The RNA-seq data from clinical prostate cancer patients’ samples were retrieved from the TCGA database (TCGA-PRAD project) and cBioportal database (SU2C prostate cancer project) using the R packages TCGAbiolinks and cBioPortalData. In total, 52 prostate normal tissues, 501 primary prostate cancer tissues, and 266 metastatic CRPC (mCRPC) tissues have been included for later analysis. Log2-transformed FPKM values plus pseudo-number 1 (log2(FPKM + 1)) were utilized for expression analysis. Moreover, the correlation between the genes was analyzed using the ggcor package.

To assess the effects of Mediator kinase inhibition on the survival of clinical PCa patients, we created a signature set of 266 genes (Table S4) that are affected by the Mediator Kinase inhibition (MKI) and inactivation in the Rv1 CRPC xenograft tumors (among the 315 DEGs listed in Table S3) and were expressed in the RNA-Seq data of clinical samples from the primary PCa patients (n= 497) and mCRPC patients (n=81) in TCGA-PRAD and SU2C projects where overall survival data were available. The expression of the 266 genes was first transformed into z-scores and the signature score for each patient was calculated by subtracting the cumulative z-scores of the genes upregulated by MKI from the cumulative z-scores of the genes downregulated by MKI. The correlation of the signature score with the overall survival among primary PCa and mCRPC patients was assessed by the Kaplan-Meier analysis using the Survminer package. To ensure accuracy, the cut-off value for the Kaplan-Meier plots were selected based on the lowest p-value within the interquartile range (25th to 75th percentile) of the gene signature scores among the patient cohort.

### Machine learning based H&E analysis

Tumor samples harvested from mice were washed with PBS and fixed in 10% neutralized paraformaldehyde for 24-38 hours. The samples were then washed with PBS twice and stored in 70% ethanol at 4°C. Fixed tumors were embedded in paraffin and sectioned at 5 μm. H&E staining was performed using standard procedures. The H&E-stained tumor sections were imaged using Leica DMIRE fluorescence microscope, controlled via Inscoper control block with Inscoper 6.3.0 software package. The images were collected using 10x Phase objective and Zeiss AxioCam MRC color camera. Acquisition of singe images (tiles) was conducted using Marzhauser Wetzlar SCAN IM 120 x 100 XY stage. Single images were stitched and corrected for brightness and contrast levels using custom ImageJ scripts. The machine learning platform WEKA v.3.3.2 was utilized to differentiate the proliferation, apoptosis and necrosis areas based on characteristic morphologies. The Artificial Intelligence (AI) models were created using manually selected reference sets of images representing proliferation, apoptosis, and necrosis, using rounds of reference data sets optimization, training of AI-models, and accurate validation of the AI-based automatic image segmentation using test images not included into the training subsets. Finally, custom ImageJ scripts were used to apply the developed AI model to whole tissue sections. The results of AI-based detection of characteristic phenotypes within all tissue sections studied were confirmed by experienced pathology experts. For visualization purposes, areas assigned to proliferation, apoptosis, and necrosis were marked with green, yellow, and red colors. Quantification of the detected areas was done using WEKA-generated probability maps and the customized ImageJ thresholding tool. All preliminary calculations were done using Precision 5820 station: Intel® Core™ i9, 18 cores, 36 threads, 3.00 GHz to 4.80 GHz Turbo, AMD® Radeon™ Pro WX 3200, 4 GB GDDR5, 4 mDP; 128 GB DDR4 RAM; M2 PCIe NVMe Class 40 SSD. Computationally heavy steps were performed using the 1060 TeraFLOPS computer cluster (Research Computing, USC, Columbia SC) with a current capacity of 1.4 petabytes.

### Immunofluorescence staining

Immunostaining was conducted on sections cut from formalin fixed paraffin embedded tissue blocks. The sections were deparaffinized and hydrated using the following steps: 10 min in xylene (twice), 7 min in 100% ethanol (twice), 7 min in 70% ethanol, and 5 min in water at room temperature (3 times). Antigen retrieval was achieved by boiling in water for 20 min. For Immunofluorescence analysis, deparaffinized and hydrated tumor sections were autoclaved at 121°C for 20 min in pH6 10mM sodium citrate, blocked with 5% normal donkey serum in PBS plus Tween 20 (PBST), then incubated overnight at 4°C in a 1 to 1000 dilution of rabbit anti-AR. Sections were then washed in PBST and incubated a 1 to 200 dilution of Alexafluor 555 Donkey anti-Rabbit IgG (Thermo Fisher) for 1 hour. Sections were then washed in PBST and counterstained for 20 minutes with 1 μM DAPI and mounted under coverslips with Prolong Glass (Thermo Fisher). Representative tumor areas were imaged with a 20x objective in an Agilent Cytation 5 imaging system. Tiff files were exported from the Agilent Gen5.12 software and ImageJ was used to create maximum intensity projections (MIPs).

### Statistical analysis

RNA-Seq experiments were conducted with a minimum of three biological replicates for each treatment condition. The procedures for RNA-Seq data analysis are detailed in previous sections. Slope and Pearson correlation coefficients were determined through linear regression and correlation analysis utilizing GraphPad Prism 9 software. qPCR analysis was carried out in biological triplicates, and the data are presented as the mean ± standard error of the mean (SEM). Statistical significance was evaluated using an ordinary two-way ANOVA and Tukey’s multiple comparisons test with GraphPad Prism 9 software. A two-tailed student’s t-test was performed for comparisons involving only two groups. The log-rank (Mantel-Cox) test, implemented in GraphPad Prism 9 software, was used to assess statistical significance between survival curves in KM analysis.

## Acknowledgements

We thank Drs. J. Chuck Harrell and Amy Olex (Virginia Commonwealth University) for advice on the separation of human and mouse RNA-Seq data in xenograft samples. We acknowledge the support of the Functional Genomics Core, Microscopy and Flow Cytometry Core and Drug Design and Synthesis Core of the COBRE Center for Targeted Therapeutics at the University of South Carolina (supported by NIH COBRE grant P20GM109091), Hollings Cancer Center and Shared Resources at the Medical University of South Carolina (partly supported by NIH grant P30 CA138313) and Chao Family Comprehensive Cancer Center Experimental Tissue Resource Shared Resource at the University of California Irvine (supported by NIH grant P30CA062203). This work was supported by NIH grants R44CA203184 (M.C., I.B.R.), R43CA203184 (D.C.P., I.B.R.), R43CA271996 (M.C., E.V.B.), R01CA266027 (I.B.R., E.V.B.,) R01CA260351 (X.Z., M.B.L.) and P20GM109091 (E.V.B., C.M., I.B.R.), Department of Defense Prostate Cancer Research Program awards W81XWH-22-1-0205 (G.W., M.C.) and W81XWH-19-1-0726 (X. Z., M.B.L.), and Support to Promote Advancement of Research and Creativity Graduate Research Grants, University of South Carolina (J.L., L.Z., X.D.).

## Author Contributions

M.C. and I.B.R. conceived the project. M.C., I.B.R, M.B.L., C.M., X.Z., D.C.P. and E.V.B. oversaw the project, designed experiments, and interpreted data. M.C., I.B.R. and J.L. wrote the manuscript. G.W., E.V.B. and B.G. reviewed and edited the manuscript. J.L., T.H., M.C., Y.L., L.W., J.L., V.S., H.J., L.Z., C.C., X.D., K.R.K., C.E.D., G.P.S., C.L., A.C. performed the experiments. M.C., J.L., H.J. and B.G. carried out bioinformatic analysis.

