## Supplemental Figures S1-S7 for "CDK19 and CDK8 Mediator kinases drive androgen-independent *in vivo* growth of castration-resistant prostate cancer"

### Figure S1

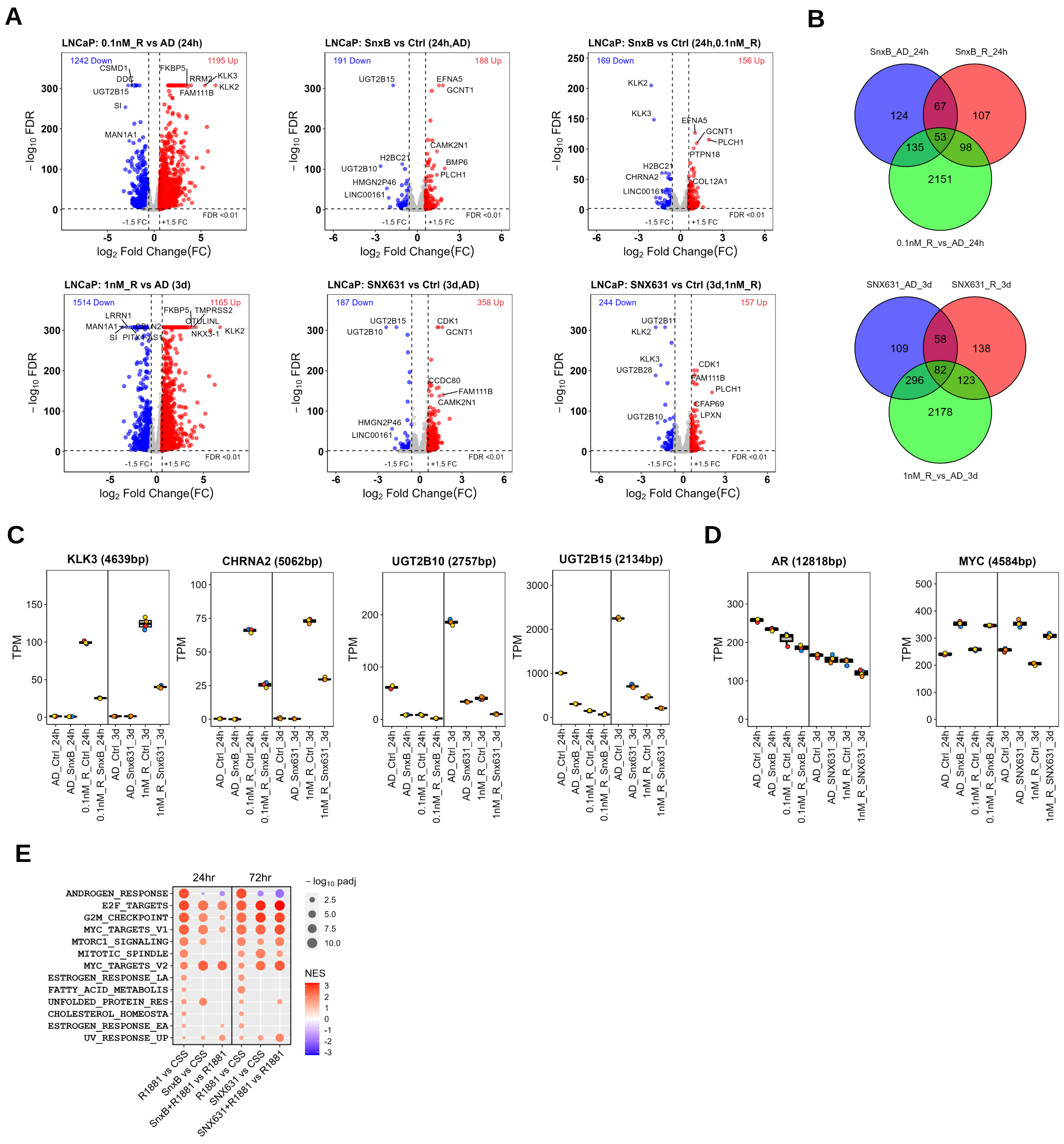

**Figure S1. RNA-Seq analysis of the effects of CDK8/19 inhibition on gene expression in LNCaP cells *in vitro*.**

**(A)** Effects of androgen stimulation and CDK8/19 inhibition by Senexin B or SNX631 under androgen-deprived and -stimulated conditions in LNCaP cells. Differentially expressed genes (DEGs) passing the selection criteria (FC >1.5, FDR <0.01) are marked with red (upregulated) and blue (downregulated) dots. **(B)** Overlap between DEGs affected by CDK8/19s and androgen stimulation. **(C)** RNA expression of the indicated genes across different conditions. **(D)** The same for AR and MYC genes. **(E)**. Hallmark pathways (GSEA) affected by androgen in LNCaP cells under the indicated conditions.

#### Figure S2

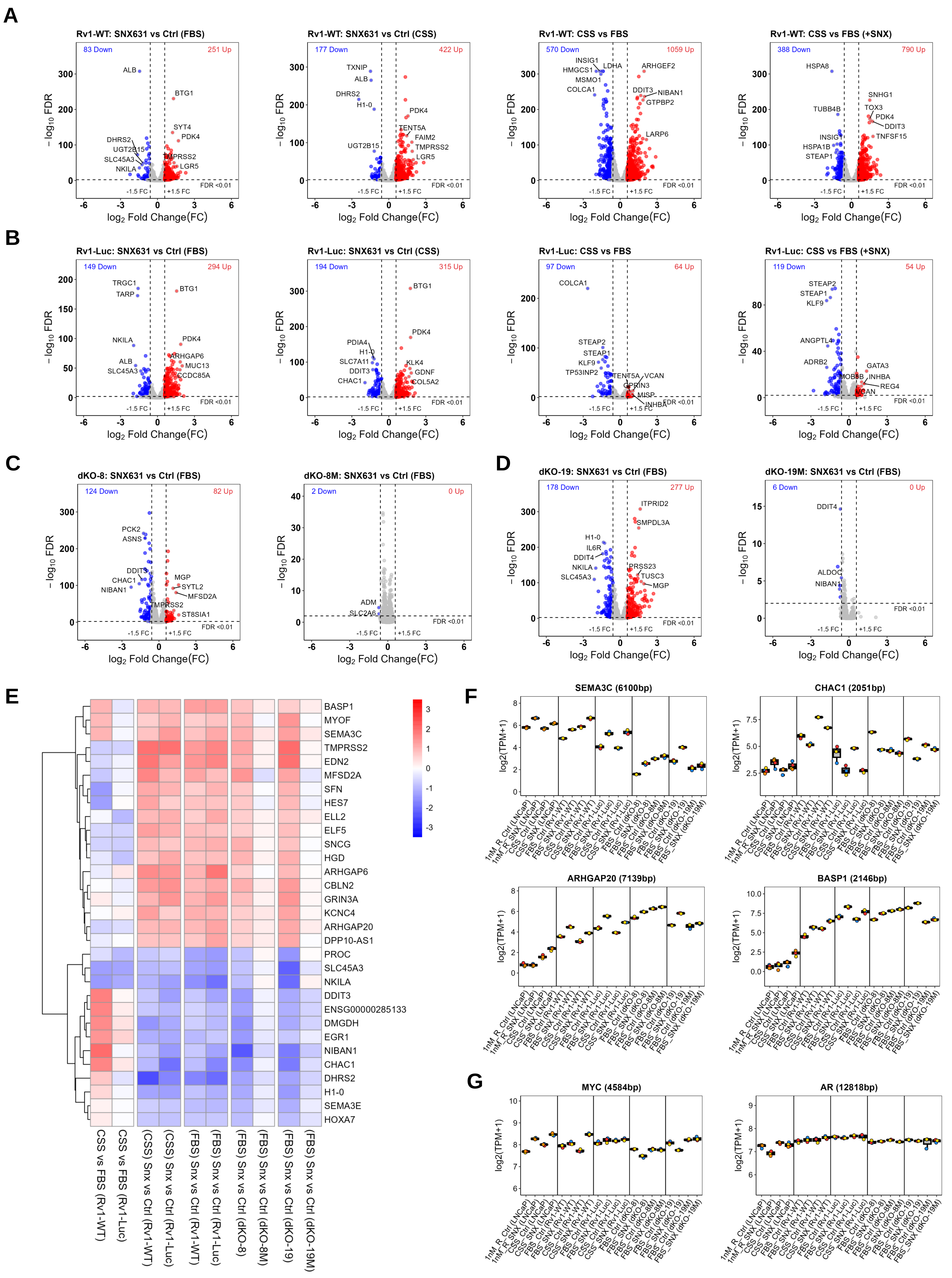
A
Rv1-WT: SNX631 vs Ctrl (FBS)
83 Down
251 Up
Rv1-WT: SNX631 vs Ctrl (CSS)
177 Down
422 Up
Rv1-WT: CSS vs FBS
570 Down
1059 Up
Rv1-WT: CSS vs FBS (+SNX)
388 Down
790 Up
B
Rv1-Luc: SNX631 vs Ctrl (FBS)
149 Down
294 Up
Rv1-Luc: SNX631 vs Ctrl (CSS)
194 Down
315 Up
Rv1-Luc: CSS vs FBS
97 Down
64 Up
Rv1-Luc: CSS vs FBS (+SNX)
119 Down
54 Up
C
dKO-8: SNX631 vs Ctrl (FBS)
124 Down
82 Up
dKO-8M: SNX631 vs Ctrl (FBS)
2 Down
0 Up
D
dKO-19: SNX631 vs Ctrl (FBS)
178 Down
277 Up
dKO-19M: SNX631 vs Ctrl (FBS)
6 Down
0 Up
log2 Fold Change(FC)
− log10 FDR
-1.5 FC
+1.5 FC
FDR <0.01
E
BASP1
MYOF
SEMA3C
TMPRSS2
EDN2
MFSD2A
SFN
HES7
ELL2
ELF5
SNCG
HGD
ARHGAP6
CBLN2
GRIN3A
KCNC4
ARHGAP20
DPP10-AS1
PROC
SLC45A3
NKILA
DDIT3
ENSG00000285133
DMGDH
EGR1
NIBAN1
CHAC1
DHRS2
H1-0
SEMA3E
HOXA7
CSS vs FBS (Rv1-WT)
CSS vs FBS (Rv1-Luc)
(CSS) Snx vs Ctrl (Rv1-WT)
(CSS) Snx vs Ctrl (Rv1-Luc)
(FBS) Snx vs Ctrl (Rv1-WT)
(FBS) Snx vs Ctrl (Rv1-Luc)
(FBS) Snx vs Ctrl (dKO-8)
(FBS) Snx vs Ctrl (dKO-8M)
(FBS) Snx vs Ctrl (dKO-19)
(FBS) Snx vs Ctrl (dKO-19M)
F
SEMA3C (6100bp)
CHAC1 (2051bp)
ARHGAP20 (7139bp)
BASP1 (2146bp)
G
MYC (4584bp)
AR (12818bp)
log2(TPM+1)

**Figure S2. RNA-Seq analysis of the effects of Mediator kinase inhibition on gene expression in 22Rv1 derivatives *in vitro*.**

**(A-B)** Effects of Mediator kinase inhibition (SNX631 vs Ctrl) under androgen-supplemented (FBS) and androgen-deprived (CSS) conditions and the effects of androgen deprivation (CSS vs FBS) in the absence or presence of SNX631 in Rv1-WT **(A)** and Rv1-Luc **(B)** cells. **(C-D)** Effects of Mediator kinase inhibition (SNX631 vs Ctrl) on dKO-8 and dKO-8M **(C)** and dKO-19 and dKO-19M **(D)** derivatives in FBS media. DEGs passing the selection criteria (FC >1.5, FDR <0.01) are marked with red (upregulated) and blue (downregulated) dots. **(E)** Heatmap of 33 DEGs commonly regulated by SNX631 in Rv1-WT, Rv1-Luc, dKO-8 and dKO-19 cells. **(F)** RNA expression of the indicated genes across different conditions. **(G)** The same for AR and MYC genes.

### Figure S3

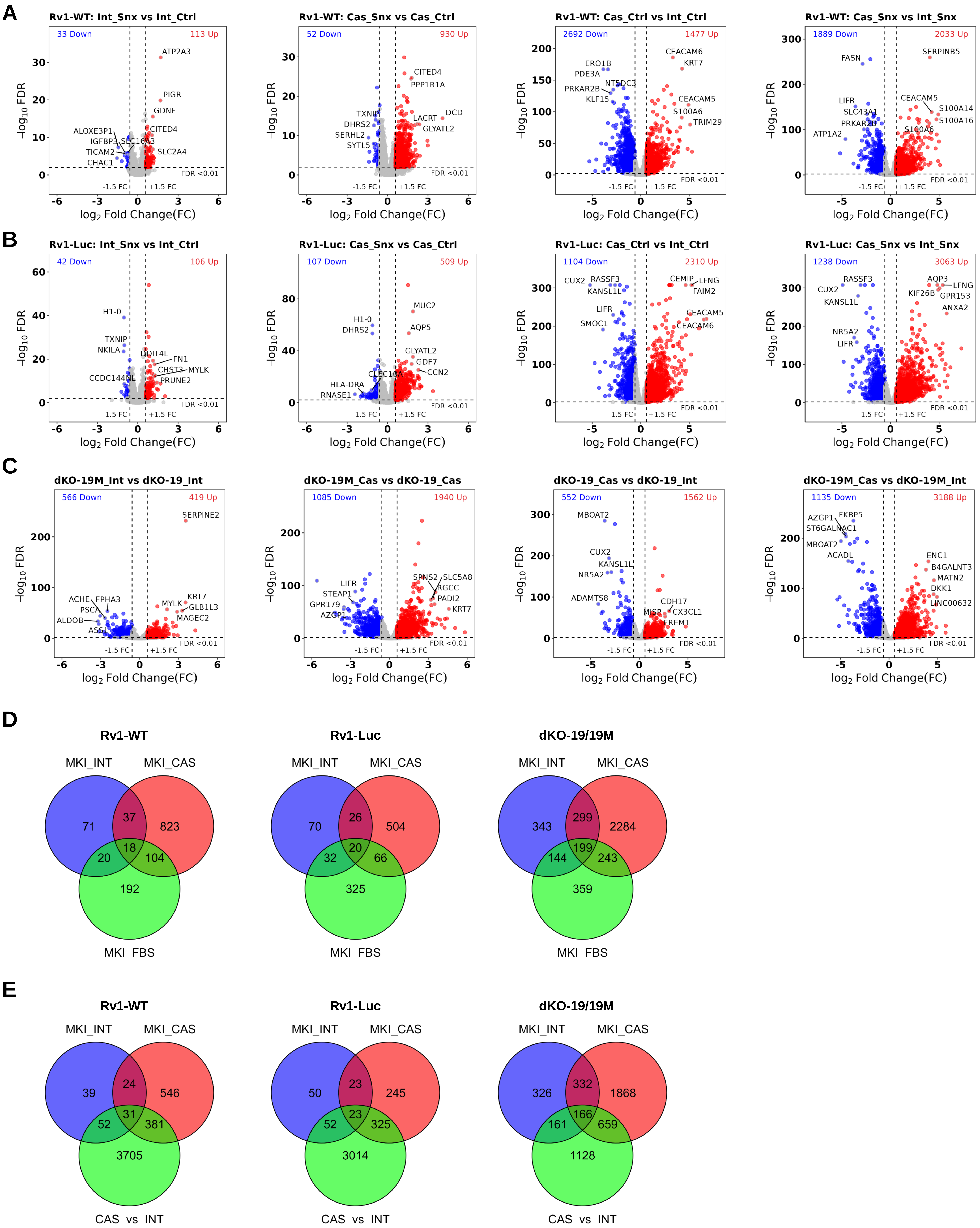
A
Rv1-WT: Int_Snx vs Int_Ctrl
33 Down
113 Up
ATP2A3
PIGR
GDNF
CITED4
ALOXE3P1
IGFBP3
SLC16A3
TICAM2
CHAC1
SLC2A4
FDR <0.01
-1.5 FC
+1.5 FC
Rv1-WT: Cas_Snx vs Cas_Ctrl
52 Down
930 Up
CITED4
PPP1R1A
TXNIP
DHRS2
SERHL2
SYTL5
LACRT
DCD
GLYATL2
Rv1-WT: Cas_Ctrl vs Int_Ctrl
2692 Down
1477 Up
ERO1B
PDE3A
NT5DC3
PRKAR2B
KLF15
CEACAM6
KRT7
CEACAM5
S100A6
TRIM29
Rv1-WT: Cas_Snx vs Int_Snx
1889 Down
2033 Up
FASN
SERPINB5
LIFR
SLC43A1
PRKAR2B
ATP1A2
CEACAM5
S100A14
S100A16
S100A6
B
Rv1-Luc: Int_Snx vs Int_Ctrl
42 Down
106 Up
H1-0
TXNIP
NKILA
DDIT4L
FN1
CHST3
MYLK
PRUNE2
CCDC144NL
Rv1-Luc: Cas_Snx vs Cas_Ctrl
107 Down
509 Up
MUC2
H1-0
DHRS2
AQP5
GLYATL2
GDF7
CCN2
CLEC10A
HLA-DRA
RNASE1
Rv1-Luc: Cas_Ctrl vs Int_Ctrl
1104 Down
2310 Up
CUX2
RASSF3
KANSL1L
CEMIP
LFNG
FAIM2
LIFR
SMOC1
CEACAM5
CEACAM6
Rv1-Luc: Cas_Snx vs Int_Snx
1238 Down
3063 Up
RASSF3
CUX2
KANSL1L
AQP3
LFNG
KIF26B
GPR153
ANXA2
NR5A2
LIFR
C
dKO-19M_Int vs dKO-19_Int
566 Down
419 Up
SERPINE2
ACHE
EPHA3
PSCA
ALDOB
ASS1
MYLK
KRT7
GLB1L3
MAGEC2
dKO-19M_Cas vs dKO-19_Cas
1085 Down
1940 Up
LIFR
STEAP1
GPR179
AZGP1
SPNS2
SLC5A8
RGCC
PADI2
KRT7
dKO-19_Cas vs dKO-19_Int
552 Down
1562 Up
MBOAT2
CUX2
KANSL1L
NR5A2
ADAMTS8
CDH17
MISP
CX3CL1
FREM1
dKO-19M_Cas vs dKO-19M_Int
1135 Down
3188 Up
AZGP1
FKBP5
ST6GALNAC1
MBOAT2
ACADL
ENC1
B4GALNT3
MATN2
DKK1
LINC00632
−log10 FDR
log2 Fold Change(FC)
D
Rv1-WT
MKI_INT
MKI_CAS
71
37
823
18
20
104
192
MKI FBS
Rv1-Luc
MKI_INT
MKI_CAS
70
26
504
20
32
66
325
MKI FBS
dKO-19/19M
MKI_INT
MKI_CAS
343
299
2284
199
144
243
359
MKI FBS
E
Rv1-WT
MKI_INT
MKI_CAS
39
24
546
31
52
381
3705
CAS vs INT
Rv1-Luc
MKI_INT
MKI_CAS
50
23
245
23
52
325
3014
CAS vs INT
dKO-19/19M
MKI_INT
MKI_CAS
326
332
1868
166
161
659
1128
CAS vs INT

**Figure S3. RNA-Seq analysis of the effects of Mediator kinase (CDK8/19) inhibition in Rv1 derivatives *in vivo* on the expression of tumor (human) genes.**

**(A-C)** Effects of Mediator kinase inhibition (SNX631 vs Ctrl or dKO-19M vs dKO-19) in intact or castrated NSG mice and the effects of castration in the absence or presence of Mediator kinase inhibition for Rv1-WT **(A)**, Rv1-Luc **(B)** and dKO-19/19M **(C)** cells *in vivo*. **(D)** Overlap of tumor DEGs affected by Mediator kinase inhibition (MKI) *in vitro* (FBS), in intact (INT) and castrated (CAS) animals for Rv1-WT, Rv1-Luc and dKO-19/19M models. **(E)** Overlap of tumor DEGs affected by Mediator kinase inhibition (MKI) or castration (CAS) in intact (INT) and castrated (CAS) animals for Rv1-WT, Rv1-Luc and dKO-19/19M models.

### Figure S4

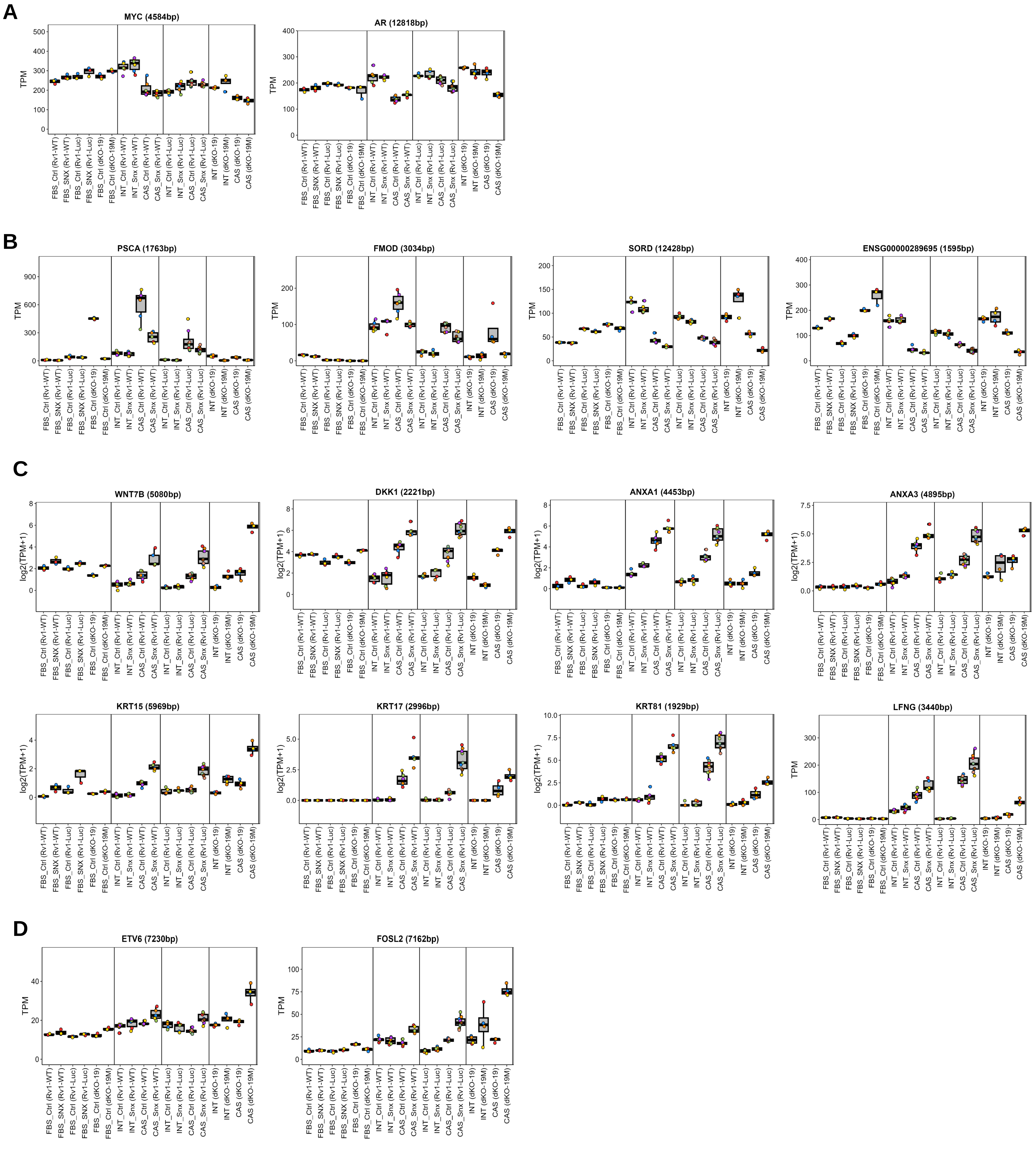

**Figure S4. Expression of representative Mediator kinase-regulated tumor (human) genes under different conditions *in vitro* and *in vivo* (RNA-Seq, TPM).**

**(A)** MYC and AR genes. **(B)** DEGs negatively regulated by Mediator kinase inhibition in 22Rv1 xenografts growing in castrated animals. **(C)** DEGs positively regulated by Mediator kinase inhibition in 22Rv1 xenografts growing in castrated animals. **(D)** ETV6 and FOSL2 genes.

### Figure S5

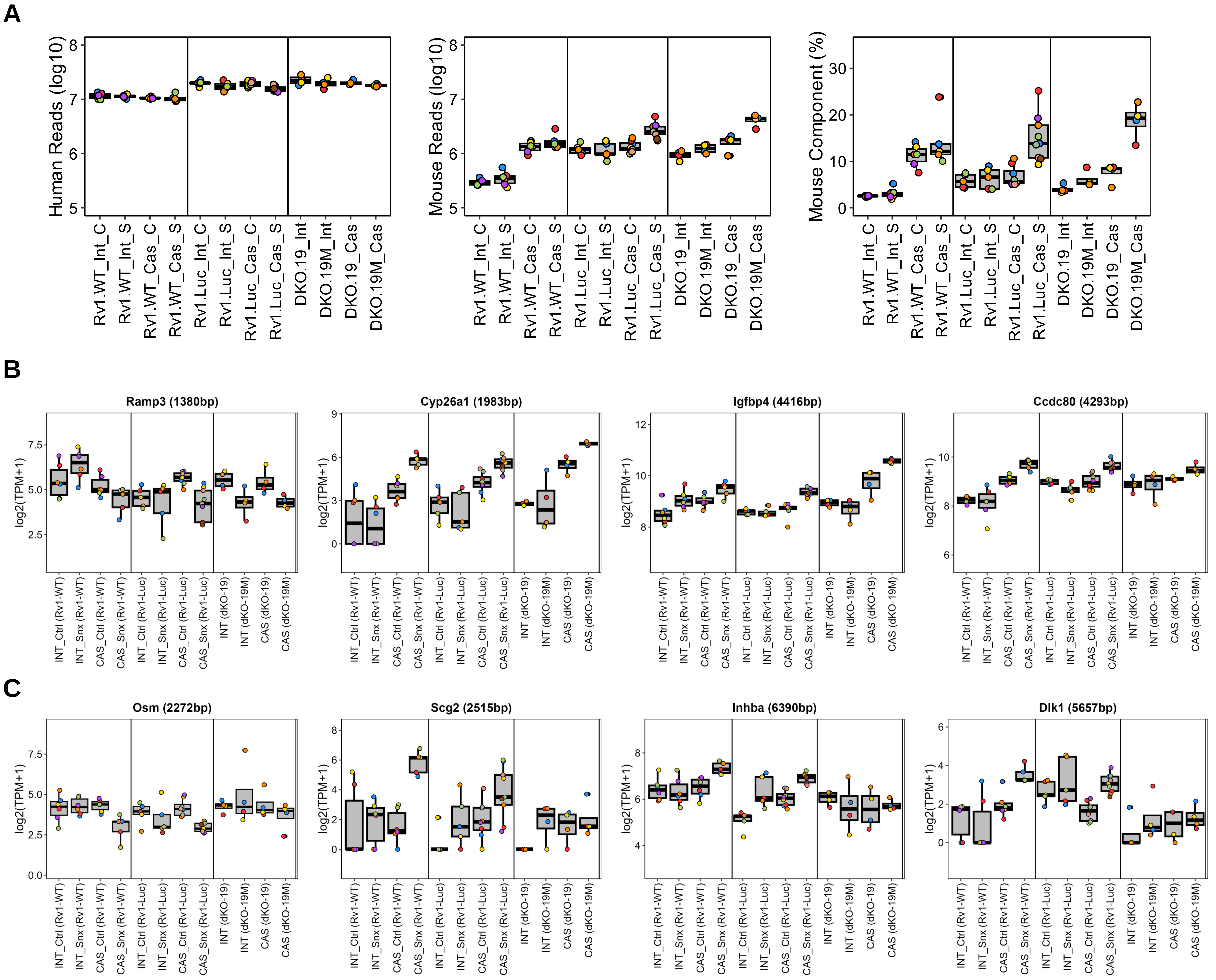

**Figure S5. RNA-Seq analysis of the effects of Mediator kinase inhibition in 22Rv1 tumors on the expression of stromal (mouse) genes.**

**(A)** Total human gene counts (left), total mouse gene counts (middle) and the percentage of mouse gene counts (right) in the indicated tumor samples. **(B,C)** Expression of representative stromal DEGs negatively or positively regulated by SNX631 treatment and Mediator kinase Mediator kinase mutagenesis **(B)** or by SNX631 treatment alone **(C)** in 22Rv1 xenografts growing in castrated animals.

### Figure S6

**A**

| PDX_ID | Tissue CEA+ | Tissue PSA+ | Prior Rx | Primary explant |
|---|---|---|---|---|
| SM0310 | Neg | Pos | Lupron, Casodex, abiraterone, docetaxel | skin metastasis |
| CG0509 | Neg | Pos | Lupron, docetaxel, carboplatin | prostatectomy |

**B**

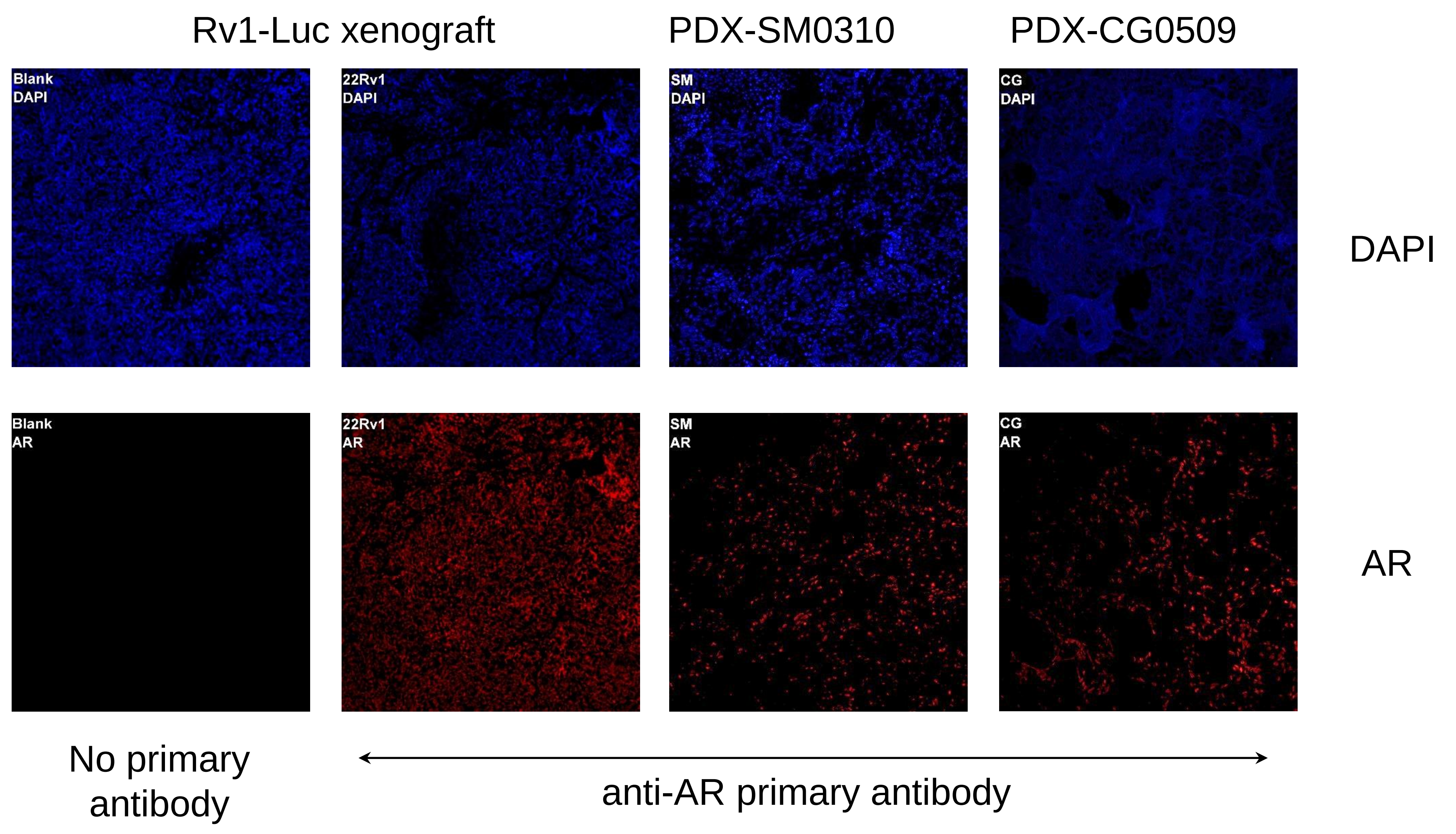

**Figure S6. Characteristics of SM0310 and CG0509 PDX models.**

**(A)** Clinical background on SM0310 and CG0509 PDX. **(B)** Immunostaining of tissue sections from Rv1-Luc, SM0310 and CG0509 xenografts for AR and DAPI.

### Figure S7

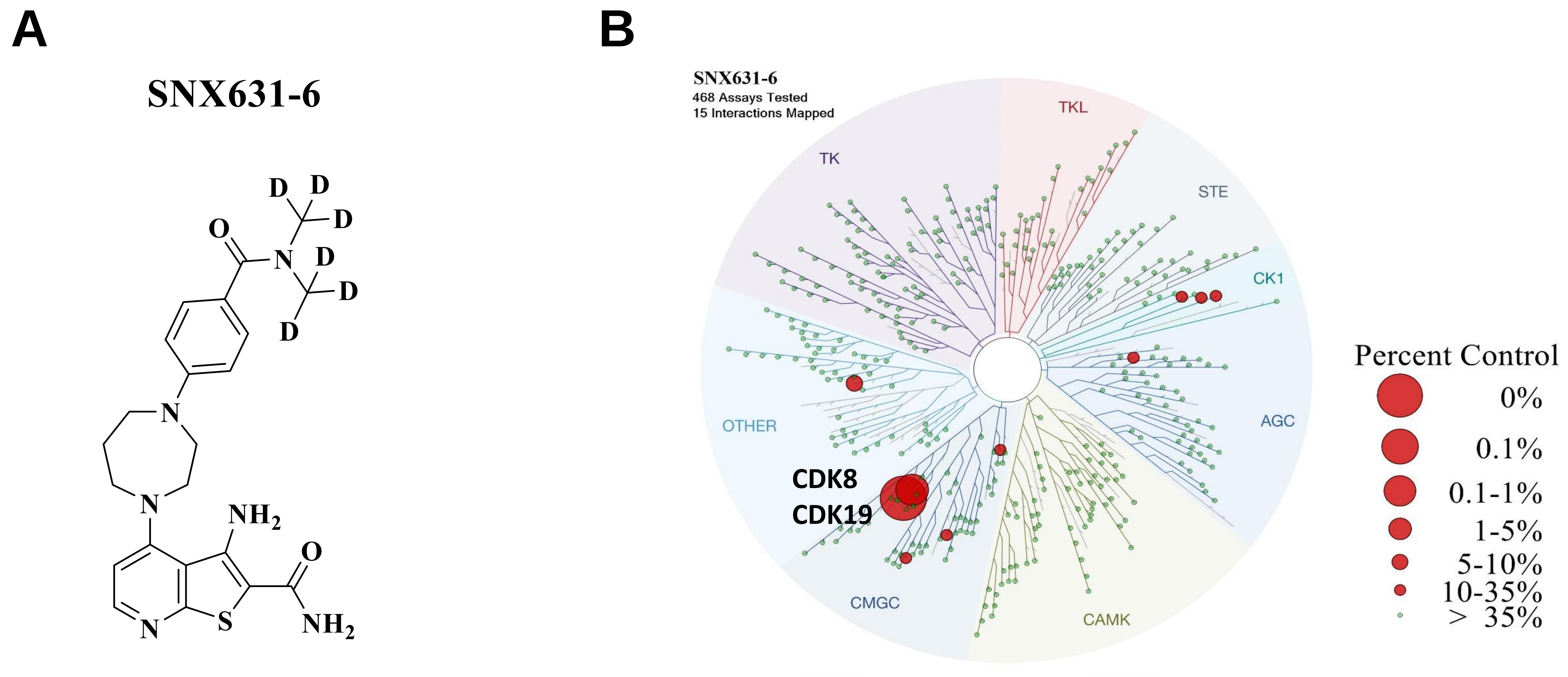

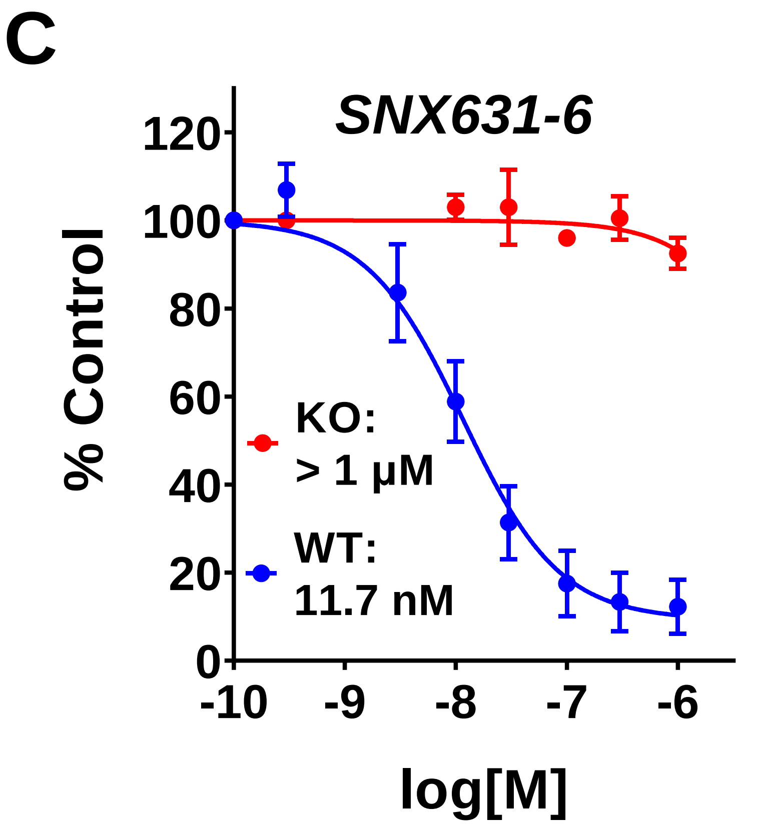

**D**

***In vitro* KINETICfinder Assay (Enzymlogic)**

| Client_ID | Target | $k_{on}$ ($M^{-1}s^{-1}$) | $k_{off}$ ($s^{-1}$) | t (min) | $T_{1/2}$ (min) | $K_d$ (M) |
|---|---|---|---|---|---|---|
| Senexin B | CDK8/cyclin C | 1.65E+06 | 3.25E-03 | 5.1 | 3.6 | 1.97E-09 |
| SNX631 | CDK8/cyclin C | 1.05E+06 | 2.83E-04 | 59 | 41 | 2.70E-10 |
| SNX631-6 | CDK8/cyclin C | 1.49E+06 | 3.72E-04 | 45 | 31 | 2.50E-10 |
| Staurosporine | CDK8/cyclin C | 1.12E+05 | 5.55E-04 | 30 | 21 | 4.95E-09 |
| Senexin B | CDK19/cyclin C | 2.13E+06 | 6.28E-03 | 2.7 | 1.8 | 2.95E-09 |
| SNX631 | CDK19/cyclin C | 1.12E+06 | 3.56E-04 | 47 | 32 | 3.19E-10 |
| SNX631-6 | CDK19/cyclin C | 9.79E+05 | 3.03E-04 | 55 | 38 | 3.09E-10 |
| Staurosporine | CDK19/cyclin C | 1.07E+05 | 2.45E-04 | 68 | 47 | 2.30E-09 |

**Figure S7. Characteristics of Mediator kinase inhibitor SNX631-6.**

**(A)** Chemical structure of SNX631-6. **(B)** Kinome profiling (Discover X) of SNX631-6 (468 kinases) at 2 µM. **(C)** Effects of SNX631-6 in a cell-based NFκB-dependent reporter assay for CDK8/19 inhibition in wildtype (WT) and CDK8/19-double knockout (KO) 293 cells. **(D)** Binding kinetics of Senexin B, SNX631, SNX631-6 and Staurosporine (a non-selective kinase inhibitor) to recombinant CDK8/CycC and CDK19/CycC proteins in the KINETICfinder Assay (Enzymlogic).
